# Sub-3 Å cryo-EM structure of RNA enabled by engineered homomeric self-assembly

**DOI:** 10.1101/2021.08.11.455951

**Authors:** Di Liu, François A. Thélot, Joseph A. Piccirilli, Maofu Liao, Peng Yin

## Abstract

Many functional RNAs fold into intricate and precise 3D architectures, and high-resolution structures are required to understand their underlying mechanistic principles. However, RNA structural determination is difficult. Herein, we present a nanoarchitectural strategy to enable the efficient single-particle cryogenic electron microscopy (cryo-EM) analysis of RNA-only structures. This strategy, termed <u>R</u>NA <u>o</u>ligomerization-enabled <u>c</u>ryo-EM via installing <u>k</u>issing-loops (ROCK), involves the engineering of target RNAs by installing kissing-loop sequences onto functionally nonessential stems for the assembly into closed homomeric nanoarchitectures. Assembly with geometric restraints leads to (1) molecular weight multiplication and (2) structural flexibility mitigation, both beneficial for cryo-EM analysis. Together with construct optimization and symmetry-expansion reconstruction, ROCK yields the cryo-EM reconstruction of the *Tetrahymena* group I intron at an overall resolution of 2.98 Å (2.85 Å resolution for the core domains), enabling the de novo model building of the complete intron RNA including previously unknown peripheral domains. When applied to smaller RNAs, ROCK readily produces modest-resolution maps, revealing the conformational rearrangement of the *Azoarcus* group I intron and the bound ligand in the FMN riboswitch. Our work unleashes the largely unexplored potential of cryo-EM in RNA structural studies.

---

In recent decades, there has been growing appreciation for the versatile and far-reaching roles that RNAs can play^1^. Besides guiding protein biosynthesis, RNAs also regulate gene expression and modulate other important biological processes by various mechanisms^2,3^, such as binding proteins, recognizing metabolites and catalyzing chemical transformations. More than 85% of the human genome is transcribed, but less than 3% of genome sequence is protein-coding^4^, suggesting a large portion of transcribed RNAs with functions and underlying structures unknown. Furthermore, the functional capacity of RNAs has been substantially expanded with *in vitro* selection and evolution^5–7^. In order to elucidate the mechanisms underlying these functional RNAs, high-resolution structural information is essential. However, experimentally solved 3D RNA structures are scarce: among the total of ~180,000 structures presently deposited with the Protein Data Bank (PDB), only 0.9% are RNA structures. This reflects the difficulties associated with the structural acquisition of RNAs, especially for those larger than 100 nucleotides (nt). RNA structural biology, traditionally being a primary territory of X-ray crystallography (and NMR for relatively small RNAs of up to ~100 nucleotides (nt)), is complicated^8^. First, the intrinsic properties of RNAs such as (a) poorly differentiated anionic surface, (b) irregular and elongated shape, and (c) structural flexibility and conformational heterogeneity, make it difficult to obtain well-diffracting crystals. Second, phase determination of RNA crystals is also difficult due to the lack of convenient and universal strategies such as selenomethionine substitution^9^ for protein crystallography.

Without the need for procuring crystals and solving the phase problem, cryo-EM is gaining increasing popularity in structural determination of protein-containing systems, and the resolution it provides is beginning to rival that of X-ray crystallography thanks to the on-going advances in instrumentation and software^10^. Nevertheless, the application of cryo-EM to RNA structural determination has not been well explored. So far, there are only two reported examples of RNA-only structures^11–13^ determined by cryo-EM that achieve a resolution of 4.5 Å or better. The first among them is the 4.5 Å structure of the *Lactococcus lactis* group IIA intron^11^, a large RNA containing > 600 nt. The other more recent case is the smaller 119-nt *Mycobacterium* sp. MCS SAM-IV riboswitch^13^: cryo-EM maps at 3.7 Å and 4.1 Å resolution were reported for the apo and ligand-bound states, respectively, but large datasets (~ two million initial particles) were required for the reconstruction. Even for the best-resolved map at 3.7 Å, the map densities for the RNA bases are barely separate and the backbone features are not well delineated, consequently making the model building process heavily dependent on computer modeling^14^.

Similar to crystallography, structural flexibility of RNA molecules is a major limiting factor for their high-resolution cryo-EM analysis: (i) double-stranded A-helix—the basic secondary structure element of RNA—has a much shorter persistence length (~64 nm, or ~230 RNA base-pairs^15^) than its protein counterpart, the α-helix (~100 nm, or~670 amino acid residues^16^); (ii) compared to proteins^17^, folded RNAs normally have fewer long-range tertiary interactions to stabilize the overall 3D architectures. In addition to flexibility, many structured RNAs are relatively small (<100 kDa, or <300 nt), making their structural determination by cryo-EM more challenging. Herein, we present ROCK (<u>R</u>NA <u>o</u>ligomerization-enabled <u>c</u>ryo-EM via installing <u>k</u>issing-loops), a nanoarchitectural engineering strategy derived from nucleic acid nanotechnology^18–22^, to address the challenges in RNA cryo-EM. Kissing-loop sequences are installed onto the peripheral stems without perturbing the functional domain to mediate self-assembly of the RNA of interest into a closed homooligomeric ring (Fig. 1a and Supplementary Figure 1). Analogous to the quaternary structures of proteins, the assembled RNA structure has a multiplied molecular weight and each constituent monomeric unit is expected to have mitigated flexibility due to the geometric restraints imposed by the kissing loops-mediated ring closure. Kissing loops, instead of sticky ends, are chosen to mediate the assembly because their paranemic characteristic^23–26^ (i.e. the two interacting loops within the kissing loops are topologically closed and can separate without the need for strand scission) minimizes the strand breaks (and thereby, the number of unique strands) and dispenses with introducing permutation.

**Figure 1.**
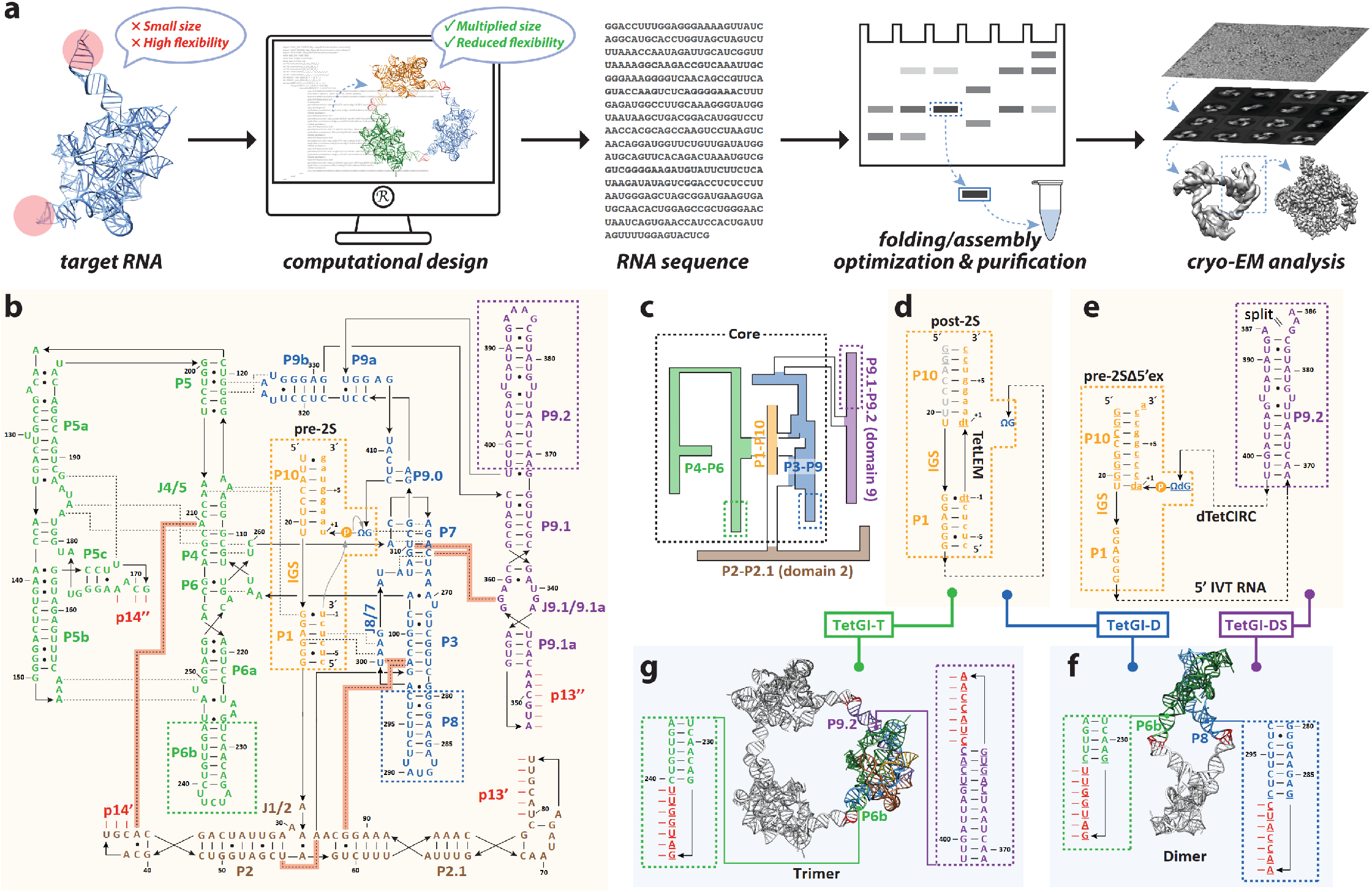
ROCK (<u>R</u>NA <u>o</u>ligomerization-enabled <u>c</u>ryo-EM via installing <u>k</u>issing-loops) and the engineering of the *Tetrahymena* group I intron (TetGI) as a case study. **a**, Workflow of ROCK. The target RNA is engineered by installing kissing-loop sequences onto the functionally nonessential, peripheral helices (highlighted by light red shadow) to form a homomeric ring (also see Supplementary Figure 1 for a more detailed comparison of the standalone RNA and assembled RNA). Native polyacrylamide gel electrophoresis (PAGE) is used to screen the optimal folding/assembly conditions and to purify the desired homooligomer containing the correctly folded RNA. The gel-purified sample is subjected to structural determination by cryo-EM. **b**, Sequence and secondary structure of the wild-type (WT) TetGI in the pre-2S state. Black arrowed lines indicate direct connections (from 5’ to 3’) in the primary sequence. Key tertiary contacts are shown with black dashed lines (newly visualized tertiary interactions revealed by this work are highlighted by red shadows). Short lines (−) indicate the canonical Watson-Crick (WC) base-pairs (bps), and dots (●) indicate the non-WC bps. ΩG is the intron’s 3’-terminal guanosine. Nucleotides involved in the P*n* (where *n* is 13 or 14) tertiary base pairing are indicated by short red sticks, and p*n*’ and p*n*” correspond to the 5’ and 3’ strands of P*n*. Lower-case nucleotides are from exons. IGS, the internal guide sequence. Dashed boxes mark the locations to be engineered. The ligation reaction catalyzed by this complex is shown by grey arrows. **c**, Schematic of the TetGI, which consists of the core domains (black dashed box; including the P4-P6, P1-P10 and P3-P9 domains), and the peripheral P2-P2.1 and P9.1-P9.2 domains (or domains 2 and 9, respectively). **d, e**, Engineered constructs corresponding to the post-2S (**d**) and the pre-2SΔ5’ex (pre-2S without 5’ exon bound; **e**) states of the intron. Underlined nucleotides are mutated from the WT. In **d**, grey nucleotides are not present due to the cleavage after U20 (see Supplementary Figure 2) during the preparation by in vitro transcription (IVT), and the ligated exon mimic (TetLEM) is chemically synthesized with two deoxy mutations. In **e**, the RNA construct has a strand split at the apical loop of P9.2 (between A386 and A387) and is formed by a 5’ RNA fragment synthesized by IVT and a chemically synthesized 3’ RNA with two deoxy mutations (dTetCIRC). **f, g**, The dimeric (**f**) and trimeric (**g**) assemblies are designed by engineering P6b and P8, and P6b and P9.2, respectively, so that the sequences (red) for forming kissing-loop motifs are installed to mediated the cohesion of the monomeric units. The design of dimer is based on the previous crystal structure^35^ of the TetGI core (PDB code: 1×8w), and the design of trimer is based on a computer model^31^ of the complete intron. In this work, three different TetGI constructs are designed and studied by cryo-EM: TetGI-D (dimeric post-2S construct), TetGI-T (trimeric post-2S construct), and TetGI-DS (dimeric pre-2SΔ5’ex construct).

Harnessing the capabilities of ROCK, we determined the structure of the complete *Tetrahymena* group I intron (TetGI)—which, as the first discovered and most iconic catalytic RNA^27^, is one of the most investigated RNA molecules and serves as a rewarding model for research of RNA biochemistry and structural biology^28,29^—at 2.98 Å resolution (2.85 Å resolution for the core). The cryo-EM map presents clear features of base-to-base separation and sugar-phosphate backbone that are characteristic for RNA, and thereby enables de novo model building of the complete TetGI, including the previously unknown peripheral domains. We also demonstrated that different regions of the TetGI can be selectively stabilized by configuring it into two different oligomers (dimer or trimer) via engineering different pairs of peripheral helices. Lastly, the generality of ROCK is validated by its application to other two smaller and more flexible RNAs—the *Azoarcus* group I intron (AzoGI; 206 nt) and FMN riboswitch (112 nt), of which modest-resolution cryo-EM maps are readily obtainable from datasets of reasonable sizes and by use of more accessible instruments. These maps allow us to capture the conformational rearrangement of the AzoGI from close to open states after the second-step of splicing, and to delineate the ligand binding environment of the FMN riboswitch, demonstrating the potential of cryo-EM for studying RNA dynamics and RNA-binding molecules.

## Results and Discussion

### Engineering the TetGI RNA for homomeric self-assembly

Figure 1b shows the secondary structure of the TetGI derived from previous prediction^30,31^ and amended by the structural study of this work. Besides the catalytic core^30^ (containing P4-P6, P3-P9 and P1-P10 domains) conserved for all group I introns, the TetGI, as a subgroup IC1 intron, also possesses peripheral insertions, P2-P2.1 and P9.1-P9.2 domains (or domains 2 and 9, respectively; Fig. 1c). Though a number of partial structures of the TetGI (including the P4-P6 domain^32,33^ and the core^34,35^) have been determined by crystallography since its discovery nearly four decades ago (see Supplementary Table 1 for a list of representative solved structures of group I introns), its complete structure has not been determined at high resolution. Nonetheless, the complete TetGI has been modeled computationally based on phylogeny, biochemistry data and a number of distance constraints that were derived from long-range tertiary interactions^30,31^ before crystal structures were available.

The TetGI catalyzes two consecutive phosphotransesterification reactions, and we chose two of its reaction states in our construct designs. Figure 1d presents a construct initially designed as the post-2S (the state after the second step of splicing) complex with two deoxy substitutions introduced at the splice junction of the ligated exon mimic (TetLEM; added in trans) to prevent the reverse reaction of the second step of splicing^36^. However, in attempting to prepare this construct, we observed that the intron RNA was cleaved co-transcriptionally between U20 and U21 as indicated by dideoxynucleotide sequencing analysis^37^ (Supplementary Figure 2), probably due to the formation of a hairpin at the 5’ end of the intron RNA. Therefore, the truncated intron RNA, when hybridized to TetLEM (Fig. 1d), can also be regarded as the complex formed by the trans-acting TetGI ribozyme (for the endonucleolytic reaction^38^) and its oligonucleotide substrate. Figure 1e shows a second construct, termed pre-2SΔ5’ex, corresponding to the pre-2S (the state before the second step of splicing) complex^36^ but without the 5’ exon. This construct is formed by two fragments: an in vitro transcribed (IVT) RNA corresponding to the 5’ fragment of the TetGI through nucleotide A386 and a chemically synthesized 37-nt chimeric oligonucleotide (dTetCIRC) corresponding to the 3’ fragment of the TetGI and the 3’ exon. To prevent the possible formation of the aforementioned hairpin presumably responsible for the cleavage near the 5’ end (Supplementary Figure 2), we introduced mutations to the 5’ sequence of the IVT RNA and compensatory mutations to the 3’ exon sequence of dTetCIRC to maintain the P10 pairing. Two deoxy substitutions are present in dTetCIRC to inhibit the hydrolysis at 3’ splice site^39^ (Supplementary Figure 3). An additional u(+1)a mutation introduces an additional base-pair at the base of P10 and potentially further improves the rigidity^36^. This pre-2SΔ5’ex complex has two features that are expected to beneficial for obtaining a higher-resolution structure: (1) compared to the post-2S complex, the extra covalent linkage between helices P9.0 and P10 may help rigidify the architecture^36^; and (2) the absence of the 5’ exon is intended to eliminate its binding/unbinding equilibrium and thereby improve the compositional and conformational homogeneity of the sample.

According to the knowledge gained from the previous structural and functional studies, stems P6b, P8 and P9.2 extend away from the catalytic core and do not participate in tertiary interactions; therefore, we chose these three stems for ROCK engineering. For the choice of kissing-loop motif throughout this work, we use the 7-bp kissing loops, which can adopt an angle from ~110° to ~130° (~120° for the average structure) based on its NMR structure^40^. Consequently, a homodimer (Fig. 1f) was designed based on the X-ray model of the core^35^ by engineering P6b and P8, and a homotrimer (Fig. 1g) was designed based on the computer model^31^ by engineering P6b and P9.2. The lengths of the stems in which the kissing-loop sequences are installed need to be optimized to ensure formation of a ring (a closed structure) instead of other linear or spiral assemblies (open structures). Promising designs expected to form rings can be readily screened in silico with software such as NanoTiler^41^ and experimentally validated by native gel electrophoresis. In this work, three TetGI constructs are designed and studied by cryo-EM: two dimeric constructs, TetGI-D and TetGI-DS, designed as the post-2S and pre-2SΔ5’ex states, respectively; and a trimeric construct, TetGI-T, designed as the post-2S state. We note that during the preparation of the paper a 6.8 Å cryo-EM structure of the TetGI in the trans-acting ribozyme form was reported^14^, and its structural information, though not used in the present study, can also guide the design of the trimeric construct.

### Sub-3 Å cryo-EM structure of the TetGI-DS construct

The best-resolved cryo-EM map of the TetGI was obtained from the TetGI-DS construct (Fig. 1e and f), so we first present the cryo-EM analysis of this construct. The characteristic shape of the TetGI-DS homodimer, clearly visible from the raw micrographs (Supplementary Figure 4) and 2D class-averages (Fig. 2a), is beneficial for particle-picking and initial alignment. Reconstruction with C_2_ symmetry applied to the homodimer delivers a cryo-EM map of a moderate resolution of 3.92 Å (see Fig. 2b for the whole C_2_ dimer, and Fig. 2c for the symmetrized monomer, or the C_2_ monomer). However, symmetry-expansion (SE) allows finer 3D classification of the monomers (Supplementary Figure 4) and refinement of these SE monomers yields a substantially improved resolution (Fig. 2d): as estimated by the Fourier shell correlation (FSC) curves (Fig. 2e), the overall resolution for the full map of the SE monomer is 2.98 Å at FSC = 0.143, with the core arriving at 2.85 Å resolution, superior to the best-resolved group I intron to date—the 3.10 Å crystal structure^36^ of the *Azoarcus* group I intron (AzoGI). This represents the first sub-3 Å cryo-EM map obtained for an RNA-only structure, enabling the *de novo* model building of the complete TetGI (Fig. 2f and g). In particular, fine details of structural features pertaining to the sugar-phosphate backbone and nucleobases of RNA are resolved (Fig. 2h; see Supplementary Figure 5 for comparing the features of different X-ray and EM maps). As a proof for the excellent map quality, we show in Fig. 2i that the density for individual bases are well separated without breaking the continuity of the backbone density at a wide range of contour levels. The distinct geometries of different types of base-pairs (bps) can be readily recognized (Fig. 2j and k), and notably, structural features of the exocyclic amino groups are visible (blue arrows in Fig. 2, i-k). Additionally, we can visualize the strong intensities of ordered Mg^2+^ ions (Fig. 2l; see Supplementary Figure 6 for the Mg^2+^ ions at the A-rich bulge of P4-P6 domain). This demonstrates the utility of cryo-EM in the localization of native metal ions in RNA structures, which have important roles in RNA folding and sometimes serve as ligands in RNA catalysis^42^. This cryo-EM approach to metal ion localization is complementary to the practice of X-ray crystallography, where heavy metals are often introduced as native metal mimics by crystal soaking to provide anomalous scattering signals.

**Figure 2.**
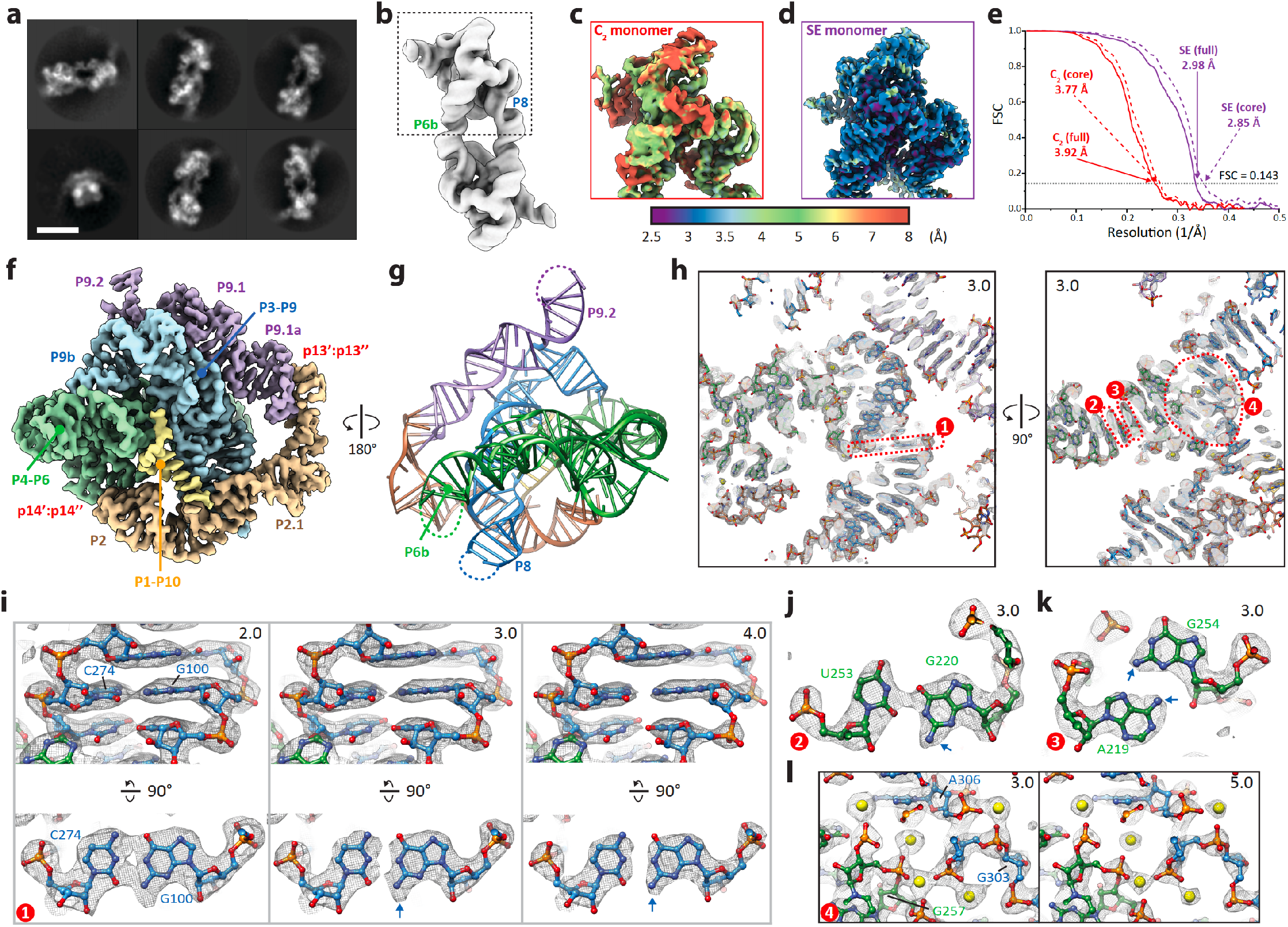
Sub-3 Å resolution cryo-EM map of TetGI-DS, a dimeric pre-2SΔ5’ex construct of the TetGI, enabled by ROCK. **a**, Representative 2D class averages of TetGI-DS dimer. Scale bar is 10 nm. **b**, Cryo-EM map of TetGI-DS dimer refined with C_2_ symmetry. **c-e**, Reconstruction with symmetry expansion (SE) improves the resolution as indicated by the local resolution maps of the monomer from the C_2_ dimer (C_2_ monomer, **c**) and the monomer after SE (SE monomer, **d**), and by the Fourier shell correlation (FSC) curves (**e**) calculated for the full (solid lines) and the core (dashed lines) of the C_2_ monomer (red) and SE monomer (purple). **f, g**, Cryo-EM map (**f**) and the derived atomic model (**g**) colored according to the secondary structure in Fig. 1b. **h**, Two clipped views of the map (grey mesh) with the atomic model (sticks for RNA and yellow spheres for Mg^2+^ ions) illustrating the quality of the high-resolution cryo-EM map. Numbers in the corners of the figure panels (through all figures in this work) indicate the root mean square deviation (RMSD, analogous to σ for X-ray crystallography maps) contour level of the raw EM map (without any zoning or carving applied unless specified otherwise). Numbered (1 through 4) red dashed boxes mark the regions in **i-l**. **i-l**, Detailed views of an example WC bp (G100:C274, **i**), a wobble bp (G220:U253, **j**), a non-canonical bp (A219:G254, **k**), and a region enclosing five Mg^2+^ ions (**l**). Blue arrows point to the feature in the map corresponding to exocyclic amino groups. In **i**, two views related by a rotation of 90° are shown with maps rendered at three different contour levels to show the map features of base-to-base separation and sugar-phosphate backbone. In **l**, the same area is shown with maps rendered at two different contour levels to show the strong map intensities of Mg^2+^ ions.

### Assembly, activity and cryo-EM analyses of TetGI-D and -T

The constructs TetGI-D and -T are both designed in the post-2S state, enabling us to investigate them in parallel to assess the validity of ROCK. Because the cation identities and concentrations can substantially influence the folding and assembly of RNA^26^, we tested the assembly of TetGI-D and -T (along with monomer control, TetGI-M) in annealing buffers containing different concentrations of Mg^2+^ and Na^+^. We then analyzed the assemblies by native polyacrylamide gel electrophoresis (PAGE) as shown in Fig. 3a. Indeed, different annealing buffers resulted in different assembly patterns of the constructs and the maximum yields of the desired dimer for TetGI-D (lanes 4 and 5) and the desired trimer for TetGI-T (lanes 10 and 11) were obtained in a buffer containing no Na^+^ and 1 or 3 mM Mg^2+^. We note that the Mg^2+^ concentrations achieving the maximum yields of the desired oligomers are close to the previously determined concentration (2 mM) optimal for the TetGI folding^43^, implying that the homomeric self-assembly system can serve as a read-out platform for optimizing conditions of the RNA folding. Consequently, we assembled the TetGI-D dimer and TetGI-T trimer in large scale with 3 mM Mg^2+^ (based on which the buffer condition for the TetGI-DS dimer assembly was chosen; see Supplementary Figure 3) and purified them by preparative native PAGE (see Methods) for the subsequent activity assays and cryo-EM analyses.

**Figure 3.**
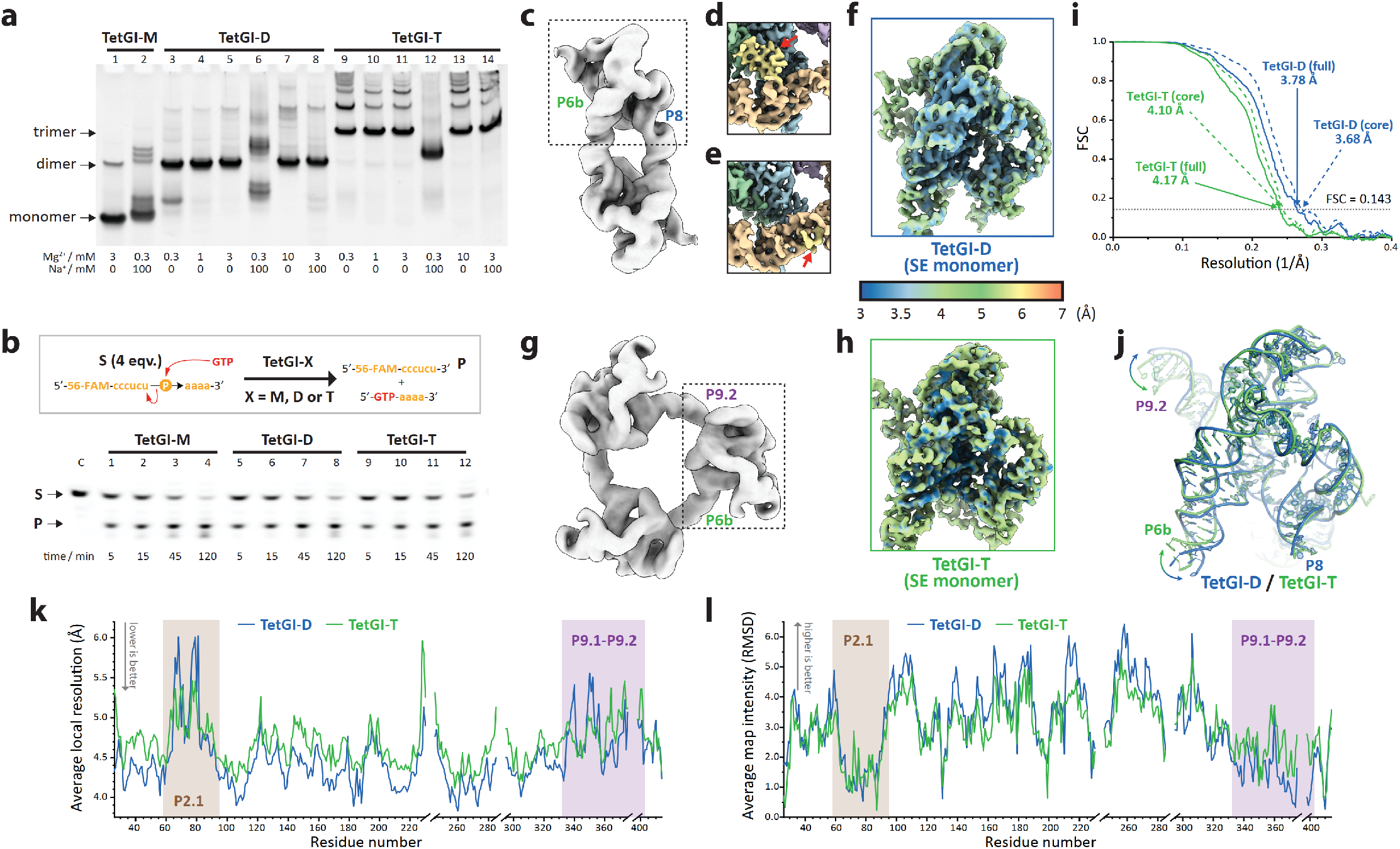
Assembly, activity and cryo-EM analyses of the dimeric and trimeric post-2S TetGI constructs, TetGI-D and -T. **a,** Native PAGE analyses of the assembly products under different annealing buffers. TetGI-M is the corresponding monomer control. The bands for the target dimer and trimer, along with the monomer control, are marked on the left of the gel image. **b**, The TetGI-D and -T constructs are catalytically active as determined by the trans-acting endonucleolytic activity. In the presence of the intron constructs (0.5 μM of monomer unit), the substrate (S, 2 μM; that is 4 equivalents (eqv.) of intron monomer unit), a 10-nt single-stranded RNA 5’ labeled with 6-fluorescein amidite (56-FAM), is cleaved by an exogenic GTP cofactor (1 mM) at 37 °C. **c,** Cryo-EM map of TetGI-D dimer refined with C_2_ symmetry. **d, e**, 3D classification of the SE monomers of TetGI-D results in two distinct conformations with (**d**) or without (**e**) the binding of TetLEM. Red arrows point to the docked double-stranded P1 (in **d**) and the single-stranded IGS (in **e**) that interacts with the minor groove of P2.1. **f**, Local resolution map of the SE monomer of TetGI-D refined by combining the two classes in **d** and **e**. See Supplementary Figure 7 for details of the reconstruction workflow for TetGI-D. **g,** Cryo-EM map of TetGI-T trimer refined with C_3_ symmetry. **h**, Local resolution map of the SE monomer of TetGI-T. The shown map is also refined by combining the two classes corresponding to the conformations with double-stranded P1 and single-stranded IGS (Supplementary Figure 8). **i**, FSC curves calculated for the full (solid lines) and the core (dashed lines) of the TetGI-D (blue) and -T (green) SE monomer. **j**, Overlay of the cryo-EM structures of TetGI-D (blue) and -T (green). Double-headed arrows mark the structural movement of P6b and P9.2 of the two structures. **k, l**, Plots of the atom-averaged local resolution (**k**) and map intensity (**l**) calculated for each residue of TetGI-D (blue) and -T (green). Though TetGI-T has an overall lower map quality (marked by higher values in **k** and lower values in **l**) than TetGI-D, it has comparable or better map quality in the regions of P2.1 and P9.1-P9.2 (highlighted by brown and purple shades, respectively) because these peripheral domains are geometrically restrained in TetGI-T.

To ensure the catalytic activity of the engineered TetGI constructs in homomeric assemblies, we assayed their trans-acting endonucleolytic activity^38^ (Fig. 3b). The reaction kinetics for TetGI-M, -D and -T are almost identical within the first 15 minutes of reaction when less than 50% of substrate is cleaved, indicating that configuring the TetGI within the homomeric assemblies does not notably affect its activity. Interestingly, TetGI-D and -T are slightly slower in cleaving the remaining substrate than TetGI-M, probably due to the tighter binding of the product to the homooligomers (i.e. via the avidity effect), which would limit the reaction rate^44^ more significantly in the later stages of the assayed reaction.

Figure 3c shows the cryo-EM map of the C_2_ dimer of TetGI-D. SE of the C_2_ dimer allows finer 3D classification (Supplementary Figure 7), revealing two conformations that are different in P1 (Fig. 3d and e; Supplementary Figure 7): Fig. 3d is the conformation with double-stranded P1 docked between the P4-P6 and P3-P9 domains^45–47^ (Supplementary Figure 7), and Fig. 3e is the conformation without TetLEM bound, so its internal guide sequence (IGS) is single-stranded and undocked. Except for P1, the other parts of the two conformations are almost identical, so the final refinement was conducted by combining these two classes, resulting in a final cryo-EM map with a resolution (overall 3.78 Å, core 3.68 Å) better than either class (Fig. 3f and Supplementary Figure 7). The cryo-EM reconstruction of TetGI-T (Fig. 3g and h, and Supplementary Figure 8) was similarly performed, and its resolution (overall 4.17 Å, core 4.10 Å) is slightly lower than that of the TetGI-D (Fig. 3i). The overall architectures of the cryo-EM structures of TetGI-D and -T are consistent with each other, and the only differences reside in the helical directions of P6b and P9.2 due to the difference of applied geometric restraints (Fig. 3j). Because TetGI-D has a higher overall resolution than TetGI-T, the map quality of TetGI-D is better than TetGI-T in most parts of the structure as indicated by the atom-averaged local resolution (Fig. 3k) and map intensity (Fig. 3l) calculated for each residue. However, in the peripheral regions of P2.1 and P9.1-P9.2, the map quality of TetGI-T is comparable to or even better than that of TetGI-D. This reflects the reduced conformational dynamics of these regions in TetGI-T due to P9.2 being geometrically restrained in the TetGI-T trimer. This result demonstrates the effectiveness of using oligomerization to mitigate structural flexibility and suggests that different regions can be preferentially stabilized by being configured within different oligomeric constructs.

### Newly visualized interactions involving the peripheral domains of the TetGI

Our cryo-EM structures of the TetGI support the configurations of peripheral domains (P2-P2.1 and P9.1-P9.2) predicted by the decades-old computer model^31^ and corroborated by a recent modest-resolution (6.8 Å) cryo-EM structure^14^. As expected for a subgroup IC1 intron, P2-P2.1 domain connects P4-P6 and P9.1-P9.2 domains via the tertiary base-pairings of P14 and P13, respectively, and these peripheral elements constitute a pseudo-continuous belt enclosing the core (Fig. 4a). Besides the overall structural organization, the present high-resolution cryo-EM structures provide a clearer view of the structural elements involving the peripheral domains (Fig. 4, b to f). The tertiary interaction of P14 (Fig. 4b) consists of the nucleotides U43, G44 and C45 (p14’) pairing with A172, C170 and G169 (p14”), respectively. An unexpected feature of P14 is the unpaired A171, which does not pair with U43 as previously predicted^31^. Also notable is that the bulged A210 from P4, which has been suggested to destabilize the folding of the isolated P4-P6 domain^33^ and was either eliminated (ΔA210 or ΔC209, ref^33,48^) or mutated (A210G, ref^35^) in previous structural studies, participates in a base-triple with a noncanonical C41:A46 pair of P2 (inset of Fig. 4b), and this long-range interaction possibly reinforces the tertiary interaction of P14. The tertiary interaction of P13 (Fig. 4c) is a 6-bp duplex formed by the base-pairings of U75 through U80 (p13’) and A352 through A347 (p13”), and stacks coaxially between the G73:C81 pair of P2.1 and the G346:C353 pair of P9.1a, bearing a conformational resemblance to some other 6-bp kissing-loop complexes^49,50^. A 4-nt bulge consisting of A69 through A72 is present near the tip of P2.1, allowing for the bent shape at the junction of P2.1 and P13.

**Figure 4.**
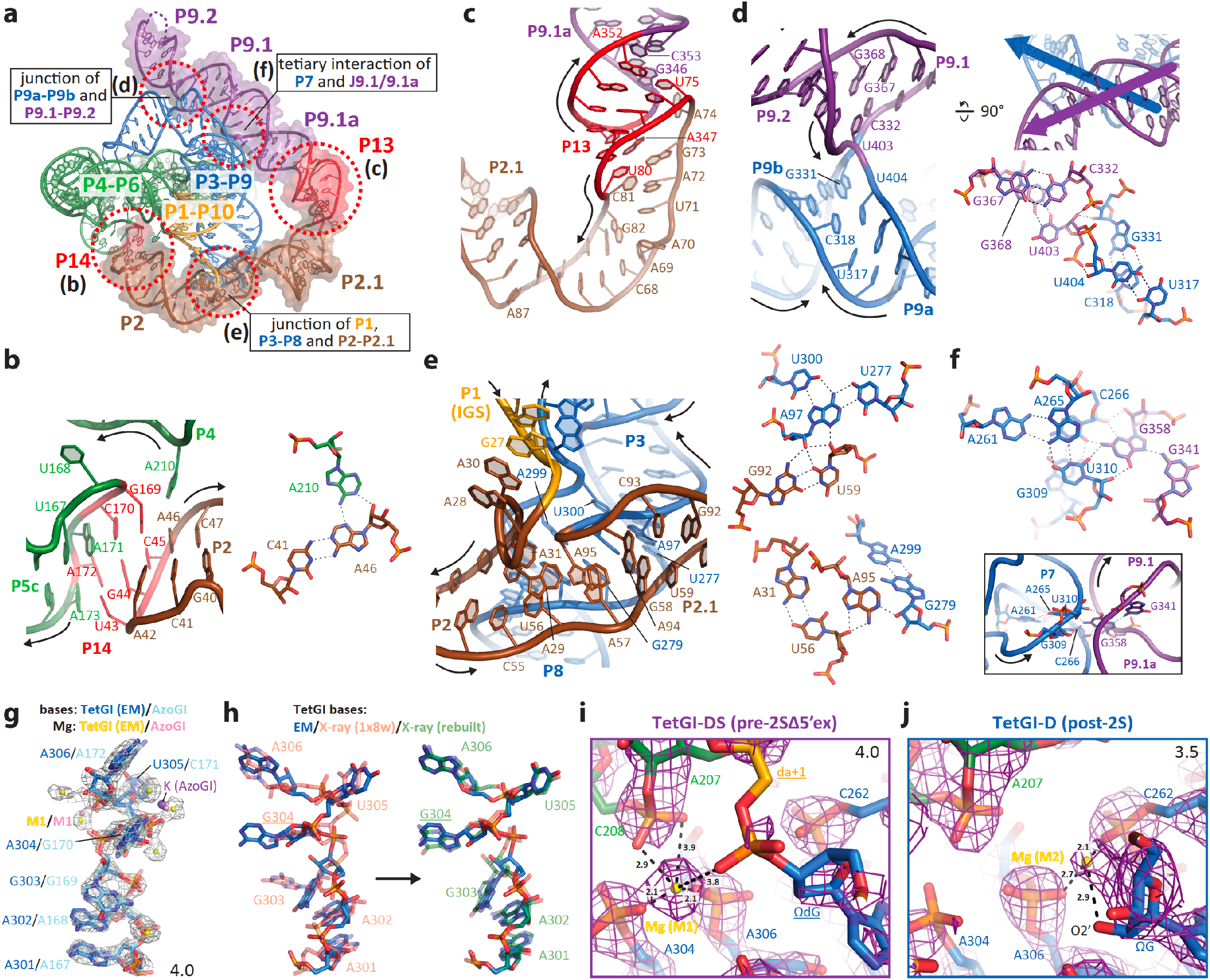
Structural insights gained from the cryo-EM structures of the TetGI. **a**, An overview of tertiary interactions and junction structures involving the peripheral domains (highlighted by surface rendering) of the TetGI. Unless specified otherwise, the TetGI structural elements are from the atomic model of TetGI-DS. **b, c**, Structural details of the tertiary interactions P14 (**b**) and P13 (**c**). Right panel of **b** shows the base-triple formed by the bulged-out A210 from P4 and the noncanonical C41:A46 pair from P2. **d, e**, Structures of the four-way junction (4WJ) at P9a-P9b and P9.1-P9.2 (**d**), and the junction at P1, P3-P8 and P2-P2.1 (**e**). Insets of **d** show the strand directions of the continuous strands of the 4WJ (top-right), and the interaction details at the crossover site (bottom-right). Top-right and bottom-right insets of **e** show the details of two tertiary contacts between the P2-P2.1 and P3-P9 domains. **f**, The tertiary contact formed by the docking of G358 of J9.1/9.1a into the minor groove of P7. **g**, A comparison of the J8/7 region of the TetGI-DS cryo-EM structure and the *Azoarcus* group I intron (AzoGI) crystal structure^36^ (PDB code: 1u6b) reveals close structural resemblance. Gray mesh is the EM map of the TetGI contoured at 4.0 RMSD level and carved within 2.0 Å of the displayed atoms. **h**, The high-resolution cryo-EM structure enables the rebuilding and re-refinement of J8/7 of the previous crystal structure of the TetGI core^35^ (PDB code: 1×8w). The structures before (light red) and after (light green) rebuilding are overlayed with the TetGI EM structure (blue). Underlined nucleotides indicate the mutations introduced for the crystallographic study. **i, j**, Active-site Mg^2+^ ions, M1 and M2, are observed in the EM maps (magenta meshes) of TetGI-DS (**i**) and TetGI-D (**j**), respectively. In the second step of splicing, M1 activates the attacking nucleophile and M2 stabilizes the leaving group. Dashed lines mark the distances (Å) between the Mg^2+^ ions and RNA heteroatoms.

Joining P9.1-P9.2 and P9a-P9b is a four-way junction (4WJ) with the flanking helices stacked in a left-handed parallel configuration^51^ (Fig. 4d and its top-right inset). Though the overall connectivity and other long-range tertiary interactions may be the major determinant for the configuration of this 4WJ, it is stabilized by the sugar-phosphate interactions of the nucleotides from the exchanging strands at the crossover site (bottom-right inset of Fig. 4d). Another covalent linkage of the core and peripheral domains is the complex multiway junction connecting P2-P2.1 with P1 and P3-P8 (Fig. 4e). Stabilizing the juxtaposition of the pseudo-continuous helices of P2-P2.1 and P3-P8 are two tertiary interactions centered by A97 and A95, respectively (insets of Fig. 4e). Within the A97-centered tertiary interaction (top-right inset of Fig. 4e), U300 forms a base-triple with A97:U277 pair, corroborating the previous biochemical evidence^52^.

Another newly visualized long-range tertiary contact between the peripheral and core domains involves the interaction of G358 from J9.1/9.1a and the minor groove of P7 (Fig. 4f), and this contact is likely to have functional significance. First, a similar contact involving the peripheral P7.2 and the minor groove of P7 also occurs in the crystal structure^53^ of the Twort group I intron (TwoGI, a subgroup IA2 intron; Supplementary Figure 9). Second, the purine-rich internal loop where the TetGI’s G358 is located is conserved in different subgroup IC1 and IE introns possessing the peripheral pseudo-continuous helix of P2.1-P13-P9.1a (Supplementary Figure 9), suggesting the presence of this contact in these introns. Because the active site of the intron is located on the major-groove side of P7, this newly visualized tertiary contact is likely to buttress the active site from the other side. Previous biochemical and chemical probing studies^54,55^ indicate that domain 9 functions to stabilize the P3-P7 region of the core and its removal affects some of the intron’s reactions involving the 3’-exon. However, previously, the major focus of the function of domain 9 has been directed to the apical loop of P9.1a (i.e. p13’’) as its participation in the P13 interaction is readily predicted by sequence complementarity^31^. Here, our high-resolution structures enabled by ROCK deliver an additional structural basis for the role of domain 9.

### J8/7 and active-site magnesium ions of TetGI

In the pre-2SΔ5’ex construct of TetGI-DS, the IGS is in the single-stranded state, propelling us to compare J8/7 (functioning as a docking site for P1) of this construct with that in the previous crystal structures of the AzoGI^36^ (with a double-stranded P1-P2; P1-P2 of the AzoGI is considered equivalent to P1 of the TetGI) and of the TetGI core^35^ (without P1 or the IGS). As shown in Fig. 4g, the configurations of J8/7 in the TetGI-DS cryo-EM structure and the AzoGI crystal structure are nearly identical. Some discrepancies lie in the bound metal ions of the two structures. For instance, we localize a Mg^2+^ ion interacting with the phosphate oxygen atoms of A301 and A302 (corresponding to A167 and A168 of the AzoGI) that has been shown theoretically to stabilize the stack-exchange junction at P3-P8^56^, but not observed in the AzoGI crystal structure^36^. As our cryo-EM model and the previous X-ray model^35^ of the TetGI differ considerably in J8/7 (Fig. 4h), we attempted to use our cryo-EM structure to rebuild and re-refine the X-ray model. Upon doing so, we observe substantial improvement of the X-ray map quality and refinement statistics (Supplementary Figure 10), and the structure differences in this region are eliminated (Fig. 4h). These results indicate that J8/7 has been pre-organized for the P1 docking, and also demonstrate the utility of high-resolution cryo-EM structures in improving the RNA model building that has been difficult for low-resolution RNA crystals.

In the cryo-EM structure of TetGI-DS, we observed an active-site Mg^2+^ (Fig. 4i) corresponding to the location of M1 observed in the crystal structure of the AzoGI ribo-ΩG pre-2S complex^57^. M1 functions as an activator of the nucleophile and the scissile phosphate in the second step of splicing^58–61^, and its presence in the 5’ exon-free intron (as in the pre-2SΔ5’ex state) suggests its possible role of activating a water molecule as the nucleophile in the 3’ splice site hydrolysis reaction^39^. However, the other metal M2 (functioning to stabilize the leaving group in the second step of splicing^62^) in the AzoGI crystal structure is not observed in our TetGI-DS construct, likely due to the deoxy substitution of ΩG (G414), which lacks the 2’-OH that coordinates to M2, ref^59^. We note that the M2 metal is present in the deoxy-ΩG pre-2S AzoGI crystal structure^36^ as a monovalent metal. In our current cryo-EM study, the only metal ion present in the buffer is Mg^2+^ (see Methods). The incapability of recruiting the divalent cation at M2 site thus is likely the cause for the suppression of the 3’ splice site hydrolysis reaction by deoxy-ΩG (Supplementary Figure 3). Nonetheless, the density for M2 could be spotted in the cryo-EM map of TetGI-D, which has a native ribo-ΩG (Fig. 4j), though we could not unambiguously build M1 due to the limited map resolution of TetGI-D. In our structures, we do not observe the density for a possible third active site metal ion that has been implicated in some functional studies^59^.

### Extending the application of ROCK to smaller RNA structures

We next set out to assess ROCK for smaller RNA structures and with more affordable, lower-performance instruments (Polara microscope with K2 camera, rather than Krios microscope with K3 camera for the TetGI structures). The AzoGI is a subgroup IC3 intron of 206 nt (~70 kDa), and compared with the TetGI, it lacks the extensive peripheral domains that facilitate RNA folding and reinforce the overall structure. Both the smaller molecular weight and the presumed increase of structure flexibility make the AzoGI a more challenging target for cryo-EM. However, the smaller size and simple fold of the AzoGI make it more attractive for crystallographic studies^36,57,63^, and the crystal structures provided the structural basis for ROCK engineering. We designed a construct, AzoGI-T, based on the post-2S complex of AzoGI by engineering its P5a and P8a for the assembly of a homotrimer (Fig. 5a and b). Similar to the TetGI-D and -T (which are also post-2S constructs; Fig. 1d), two deoxy substitutions are introduced in the in-trans added ligated exon mimic (AzoLEM). Using a similar workflow as the TetGI constructs presented earlier (Supplementary Figures 11 and 12), we obtained a 4.9 Å resolution cryo-EM map of the SE monomer (Fig. 5c and d) corresponding to a conformation with P1-P10 tightly docked (Fig. 5e), which is from the most populated and best resolved class from 3D classification (Supplementary Figure 12) and is similar to the post-2S AzoGI crystal structure^63^. Additionally, we reconstructed the cryo-EM map from another class of particles corresponding to an alternative open conformation (Fig. 5f). Though this map is of a substantially lower resolution (~ 8Å), the relative movement of the P2-P1-P10 and P4-P5 could be clearly discerned (Fig. 5f). In previous crystallographic studies^36,57,63^ of the AzoGI, such a large conformational change has not been observed among different constructs, likely due to the constraint applied by the similar crystal-packing interfaces. Thus, ROCK retains or even boosts the capability of cryo-EM in the study of functional conformational dynamics of RNA while restraining the nonfunctional structural flexibility to enable finer 3D classification.

**Figure 5.**
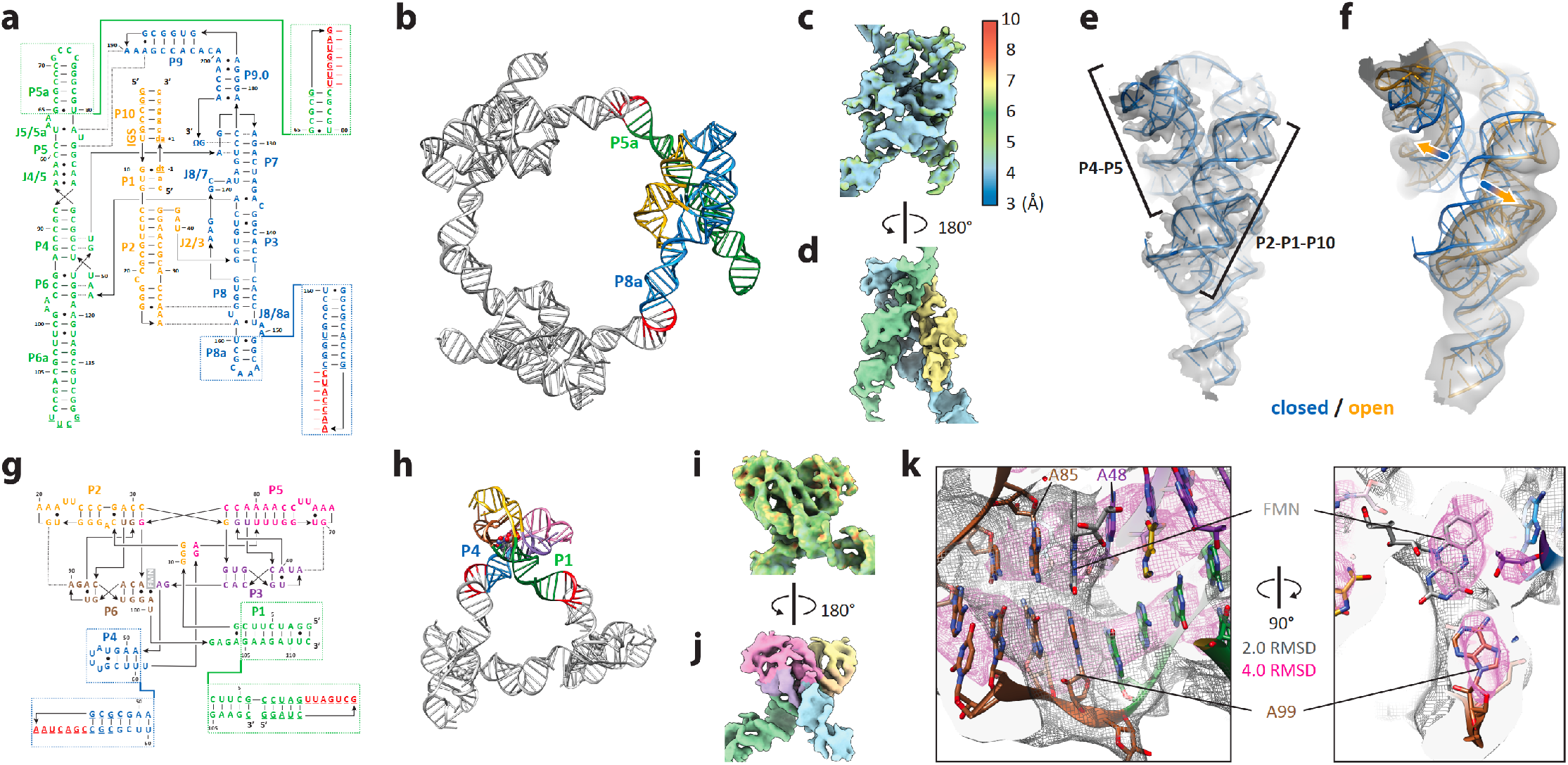
The homomeric self-assembly strategy applied to two smaller structured RNAs. **a**, Sequence and secondary structure of the AzoGI. Dashed boxes mark the engineering of P5a and P8a in the construct for trimeric self-assembly, AzoGI-T. **b**, Computer model of the assembled homotrimer of AzoGI-T. **c, d**, Cryo-EM maps rendered by local resolution (**c**) and coloring based on the secondary structure in **a** (**e**). **e**, **f,** A comparison of the refined cryo-EM maps of the best-resolved (4.9 Å; the same as the map in **c, d**) class of particles corresponding to the closed conformation (**e**) from 3D classification of the SE monomer and another less-resolved (~8.0 Å) one corresponding to the open conformation. In **f**, the fitted atomic model of the open conformation (orange) is overlayed with that of the close conformation (blue) and the arrows mark the structural movement of P2-P1-P10 and P4-P5. **g**, Sequence and secondary structure of the *Fusobacterium nucleatum* FMN riboswitch. Dashed boxes mark the engineering of P1 and P4 in the construct for trimeric self-assembly, FMNrsw-T. **h**, Computer model of the assembled homotrimer of FMNrsw-T. **i, j**, Cryo-EM maps rendered by local resolution (**i**) and coloring based on the secondary structure in **g** (**j**). The color keys for local resolution of **i** are the same as those shown in **c**. **k**, Clipped views showing the cryo-EM map of the FMN ligand and its binding environment. The cryo-EM map is rendered at two different contour levels (grey mesh at 2.0 RMSD and magenta mesh at 4.0 RMSD).

Lastly, we challenged ROCK to an even smaller target—the *Fusobacterium nucleatum* FMN riboswitch of 112 nt (~35 kDa). Based on its crystal structure^64^, we installed the kissing-loop sequences onto its P1 and P4 for the assembly of a homotrimer (Fig. 5g and h; this construct is referred to as FMNrsw-T). Interestingly, this construct assembles into two different homooligomers, dimer or trimer, when annealed in different buffer conditions (Supplementary Figure 13). Ligand-binding assays (Supplementary Figure 13) show that the RNA within the expected trimer is in its functionally relevant conformation. In contrast, the dimer is less competent in ligand binding and is likely a kinetic product. It is also possible that the monomeric subunit in the dimer is captured in an alternate conformation of the apo riboswitch^65^. The purified trimer was mixed with the FMN ligand and was then subjected to cryo-EM analysis (Supplementary Figure 14). Due to the small size of this RNA, the final refinement after SE was conducted on the whole trimer particles with one SE monomer focused (Supplementary Figure 14). Ultimately, we obtained the 5.9 Å resolution cryo-EM map of the focused monomer within the FMNrsw-T trimer (Fig. 5i and j). The bound ligand along with its binding environment can be visualized in this modest-resolution cryo-EM map due to the strong map intensities of the ligand and its vicinity (Fig. 5k). While we attribute the limited resolutions of AzoGI-T and FMNrsw-T to their smaller sizes and increased structural flexibility compared to the TetGI constructs, we could not exclude the possibility that other experimental variables, such as instruments and ice thickness, may also affect the achievable resolution.

## Conclusion

In this work, we have determined the cryo-EM structures of three different RNAs with sizes ranging from ~35 to ~140 kDa and belonging to two important categories of functional RNAs—ribozymes and riboswitches, showcasing the generality of ROCK. The core of ROCK is construct engineering for homooligomeric self-assembly, which facilitates the whole pipeline of RNA cryo-EM determination from RNA folding optimization to cryo-EM reconstruction. In fact, construct engineering is also a very common strategy employed in RNA crystallography^8^ to create preferred intermolecular crystal-packing interactions, which, besides mediating the crystal growth, also serve to dampen the structural flexibility. Compared to engineering crystal-packing interactions, designing RNA constructs for homomeric self-assembly is more tractable thanks to the programmability of RNAs and the advances of nucleic acid nanotechnology^18–22^.

It is important to note the technical requirements of construct engineering for ROCK. Firstly, the workflow of ROCK starts with construct engineering that requires the RNA have at least two nonfunctional helices, which can be usually identified in an RNA species (within a particular RNA family of interest) possessing more extensive peripheral structural elements or conferred by performing the operation of motif fusion^26^. Secondly, construct engineering can be substantially facilitated by an initial structural understanding, which can be readily obtained by solved analogous or partial structures (by crystallography or NMR), computer modeling^31^, atomic force microscopy^66^, small-angle X-ray scattering^67^, comparative gel electrophoresis^68^, and preliminary low- or modest-resolution cryo-EM models^14^. It is also important to note that, for all structural biology studies, one has to ensure the biological meaningfulness of the obtained structural insights. Specifically for ROCK, to exclude the possibility that *de novo* designed oligomerization alters the native structure and function, we assayed the activities of the engineered, assembled RNAs.

Due to the structural flexibility and folding heterogeneity that are well known to complicate the structure and function analysis of RNA, previous RNA cryo-EM studies^12–14^ aiming for high-resolution structures entail a large number of initial particles, making the workflow experimentally and computationally demanding. Our strategy greatly mitigates the nonfunctional conformational dynamics by geometric restraints and improves the sample homogeneity by natively purifying the target homooligomers. Further, the resulting symmetric assemblies are also preferred subjects for cryo-EM: (i) the characteristic shapes are more convenient for initial alignment of the particles; and (ii) special EM image processing procedures such as symmetry expansion and individual subunit-focused classification and refinement can be utilized for achieving unprecedented resolution for RNA-only structures. The principle of ROCK is also envisioned to apply for protein-containing systems: one exciting possibility is the direct engineering of small monomeric proteins for self-assembly to multiple the molecular weight and reduce structural flexibility, which differs from a recently devised approach of attaching small proteins to large homomeric scaffolds^69,70^ aiming to increase the molecular weight. In conclusion, we believe that ROCK unleashes the largely unexplored potential of cryo-EM in RNA structural studies, opening new opportunities for elucidating the mechanisms of functional RNAs and facilitating the design- and structure-based approaches to the invention of RNA-targeting therapeutics^71^.

## Data availability

The data supporting the findings of this study are principally within the figures and the associated Supplementary Information. Atomic coordinates and cryo-EM maps have been deposited with the Protein Data Bank and the Electron Microscopy Data Bank under the accession codes: 7R6L and EMD-24281 for TetGI-DS, 7R6M and EMD-24282 for TetGI-D, 7R6N and EMD-24283 for TetGI-T, EMD-24284 for AzoGI-T, and EMD-24285 for FMNrsw-T. Additional data are available from the authors upon request.

## Acknowledgments

We thank Eric Westhof, Thomas Cech, Feng Guo, Mingjie Dai and Yaming Shao for helpful discussions, Svetla Stoilova-McPhie for the help in pilot EM experiments, and Melanie Ziegler for proofreading. D. L. is a Merck Fellow of the Life Sciences Research Foundation. This work was supported by NSF grants (CMMI-1333215, CMMI-1344915 and CBET-1729397), AFOSR grant (MURI FATE, FA9550-15-1-0514), NIH grant (5DP1GM133052) and internal support from the Wyss Institute to P.Y, NIH grant (R01GM122797) to M.L., and NIH grant (R01GM102489) to J.A.P.

## Author Contributions

D.L., M.L. and P.Y. conceived the project and designed the study; D.L. designed and prepared the constructs, performed the functional assays, and conducted pilot EM experiments; F.A.T. collected the EM data; F.A.T. and M.L. performed the reconstruction and generated the maps; D.L. built and refined the atomic models with F.A.T.; D.L., F.A.T. and J.A.P. drafted the manuscript; J.A.P., M.L. and P.Y. advised the experiments; all authors analyzed the data and commented on the manuscript.

## Supplementary Information

is available with the paper.

## References

1. Breaker, R. R. & Joyce, G. F. The expanding view of RNA and DNA function. Chem. Biol. 2014, 21(9): 1059–1065.

2. Mortimer, S. A., Kidwell, M. A. & Doudna, J. A. Insights into RNA structure and function from genome-wide studies. Nat Rev Genet 2014, 15(7): 469–479.

3. Serganov, A. & Patel, D. J. Ribozymes, riboswitches and beyond: regulation of gene expression without proteins. Nat Rev Genet 2007, 8(10): 776–790.

4. Hangauer, M. J., Vaughn, I. W. & McManus, M. T. Pervasive transcription of the human genome produces thousands of previously unidentified long intergenic noncoding RNAs. PLoS Genet 2013, 9(6): e1003569.

5. Robertson, D. L. & Joyce, G. F. Selection in vitro of an RNA enzyme that specifically cleaves single-stranded DNA. Nature 1990, 344(6265): 467–468.

6. Ellington, A. D. & Szostak, J. W. In vitro selection of RNA molecules that bind specific ligands. Nature 1990, 346(6287): 818–822.

7. Tuerk, C. & Gold, L. Systematic evolution of ligands by exponential enrichment: RNA ligands to bacteriophage T4 DNA polymerase. Science 1990, 249(4968): 505–510.

8. Zhang, J. & Ferre-D’Amare, A. R. New molecular engineering approaches for crystallographic studies of large RNAs. Curr Opin Struct Biol 2014, 26: 9–15.

9. Hendrickson, W. A., Horton, J. R. & LeMaster, D. M. Selenomethionyl proteins produced for analysis by multiwavelength anomalous diffraction (MAD): a vehicle for direct determination of three-dimensional structure. The EMBO Journal 1990, 9(5): 1665–1672.

10. Nogales, E. The development of cryo-EM into a mainstream structural biology technique. Nat Methods 2016, 13(1): 24–27.

11. Qu, G. et al. Structure of a group II intron in complex with its reverse transcriptase. Nat Struct Mol Biol 2016, 23(6): 549–557.

12. Li, S. et al. Structural basis of amino acid surveillance by higher-order tRNA-mRNA interactions. Nat Struct Mol Biol 2019, 26(12): 1094–1105.

13. Zhang, K. et al. Cryo-EM structure of a 40 kDa SAM-IV riboswitch RNA at 3.7 A resolution. Nat Commun 2019, 10(1): 5511.

14. Kappel, K. et al. Accelerated cryo-EM-guided determination of three-dimensional RNA-only structures. Nat Methods 2020, 17(7): 699–707.

15. Abels, J. A., Moreno-Herrero, F., van der Heijden, T., Dekker, C. & Dekker, N. H. Single-molecule measurements of the persistence length of double-stranded RNA. Biophys. J. 2005, 88(4): 2737–2744.

16. Choe, S. & Sun, S. X. The elasticity of alpha-helices. J Chem Phys 2005, 122(24): 244912.

17. Kessel, A. & Ben-Tal, N. Introduction to proteins: structure, function, and motion. CRC Press, 2010.

18. Seeman, N.C. Nanomaterials based on DNA. Annual Review of Biochemistry 2010, 79: 65–87.

19. Seeman, N.C. Structural DNA Nanotechnology. Cambridge University Press, 2016.

20. Seeman, N.C. & Sleiman, H. F. DNA nanotechnology. Nature Reviews Materials 2017, 3(1): 17068.

21. Guo, P. The emerging field of RNA nanotechnology. Nature Nanotechnol. 2010, 5(12): 833–842.

22. Grabow, W. W. & Jaeger, L. RNA self-assembly and RNA nanotechnology. Acc Chem Res 2014, 47(6): 1871–1880.

23. Zhang, X., Yan, H., Shen, Z. & Seeman, N. C. Paranemic Cohesion of Topologically-Closed DNA Molecules. Journal of the American Chemical Society 2002, 124(44): 12940–12941.

24. Wang, X. et al. Paranemic Crossover DNA: There and Back Again. Chem Rev 2018, 119(10): 6273–6289.

25. Geary, C. W. & Andersen, E. S. Design Principles for Single-Stranded RNA Origami Structures. 2014; Cham: Springer International Publishing; 2014. p. 1–19.

26. Liu, D. et al. Branched kissing loops for the construction of diverse RNA homooligomeric nanostructures. Nature Chem. 2020, 12(3): 249–259.

27. Kruger, K. et al. Self-splicing RNA: Autoexcision and autocyclization of the ribosomal RNA intervening sequence of tetrahymena. Cell 1982, 31(1): 147–157.

28. Hougland, J. L., Piccirilli, J. A., Forconi, M., Lee, J. & Herschlag, D. How the Group I Intron Works: A Case Study of RNA Structure and Function. In: Gesteland RF, Atkins JF, Cech TR (eds). RNA World, 3rd edn. Cold Spring Harbor Laboratory Press, 2006, pp 133–205.

29. Golden, B. L. Group I Introns: Biochemical and Crystallographic Characterization of the Active Site Structure. Ribozymes and RNA Catalysis. The Royal Society of Chemistry, 2007, pp 178–200.

30. Michel, F. & Westhof, E. Modelling of the three-dimensional architecture of group I catalytic introns based on comparative sequence analysis. Journal of Molecular Biology 1990, 216(3): 585–610.

31. Lehnert, V., Jaeger, L., Michel, F. & Westhof, E. New loop-loop tertiary interactions in self-splicing introns of subgroup IC and ID: a complete 3D model of the Tetrahymena thermophila ribozyme. Chemistry & Biology 1996, 3(12): 993–1009.

32. Cate, J. H. et al. Crystal structure of a group I ribozyme domain: principles of RNA packing. Science 1996, 273(5282): 1678–1685.

33. Juneau, K., Podell, E., Harrington, D. J. & Cech, T. R. Structural Basis of the Enhanced Stability of a Mutant Ribozyme Domain and a Detailed View of RNA–Solvent Interactions. Structure 2001, 9(3): 221–231.

34. Golden, B. L., Gooding, A. R., Podell, E. R. & Cech, T. R. A preorganized active site in the crystal structure of the Tetrahymena ribozyme. Science 1998, 282(5387): 259–264.

35. Guo, F., Gooding, A. R. & Cech, T. R. Structure of the Tetrahymena ribozyme: base triple sandwich and metal ion at the active site. Mol Cell 2004, 16(3): 351–362.

36. Adams, P. L., Stahley, M. R., Kosek, A. B., Wang, J. & Strobel, S. A. Crystal structure of a self-splicing group I intron with both exons. Nature 2004, 430(6995): 45–50.

37. Hahn, C. S., Strauss, E. G. & Strauss, J. H. Dideoxy sequencing of RNA using reverse transcriptase. Methods Enzymol. 1989, 180: 121–130.

38. Zaug, A. J., Been, M. D. & Cech, T. R. The Tetrahymena ribozyme acts like an RNA restriction endonuclease. Nature 1986, 324(6096): 429–433.

39. Inoue, T., Sullivan, F. X. & Cech, T. R. New reactions of the ribosomal RNA precursor of Tetrahymena and the mechanism of self-splicing. Journal of Molecular Biology 1986, 189(1): 143–165.

40. Lee, A. J. & Crothers, D. M. The solution structure of an RNA loop–loop complex: the ColE1 inverted loop sequence. Structure 1998, 6(8): 993–1007.

41. Bindewald, E., Grunewald, C., Boyle, B., O’Connor, M. & Shapiro, B. A. Computational strategies for the automated design of RNA nanoscale structures from building blocks using NanoTiler. J Mol Graph Model 2008, 27(3): 299–308.

42. Woodson, S. A. Metal ions and RNA folding: a highly charged topic with a dynamic future. Curr Opin Chem Biol 2005, 9(2): 104–109.

43. Rook, M. S., Treiber, D. K. & Williamson, J. R. An optimal Mg(2+) concentration for kinetic folding of the tetrahymena ribozyme. Proc. Natl. Acad. Sci. USA 1999, 96(22): 12471–12476.

44. Herschlag, D. & Cech, T. R. Catalysis of RNA cleavage by the Tetrahymena thermophila ribozyme. 2. Kinetic description of the reaction of an RNA substrate that forms a mismatch at the active site. Biochemistry 1990, 29(44): 10172–10180.

45. Pyle, A. M. & Cech, T. R. Ribozyme recognition of RNA by tertiary interactions with specific ribose 2’-OH groups. Nature 1991, 350(6319): 628–631.

46. Herschlag, D., Eckstein, F. & Cech, T. R. Contributions of 2’-hydroxyl groups of the RNA substrate to binding and catalysis by the Tetrahymena ribozyme. An energetic picture of an active site composed of RNA. Biochemistry 1993, 32(32): 8299–8311.

47. Strobel, S. A. & Cech, T. R. Tertiary interactions with the internal guide sequence mediate docking of the P1 helix into the catalytic core of the Tetrahymena ribozyme. Biochemistry 1993, 32(49): 13593–13604.

48. Ye, J. D. et al. Synthetic antibodies for specific recognition and crystallization of structured RNA. Proc. Natl. Acad. Sci. USA 2008, 105(1): 82–87.

49. Ennifar, E., Walter, P., Ehresmann, B., Ehresmann, C. & Dumas, P. Crystal structures of coaxially stacked kissing complexes of the HIV-1 RNA dimerization initiation site. Nature Struct. Biol. 2001, 8(12): 1064–1068.

50. Lebars, I. et al. Exploring TAR-RNA aptamer loop-loop interaction by X-ray crystallography, UV spectroscopy and surface plasmon resonance. Nucleic Acids Res. 2008, 36(22): 7146–7156.

51. Lilley, D. M. Structures of helical junctions in nucleic acids. Quarterly Reviews of Biophysics 2000, 33(2): 109–159.

52. Szewczak, A. A. et al. An important base triple anchors the substrate helix recognition surface within the Tetrahymena ribozyme active site. Proc. Natl. Acad. Sci. USA 1999, 96(20): 11183–11188.

53. Golden, B. L., Kim, H. & Chase, E. Crystal structure of a phage Twort group I ribozyme-product complex. Nat Struct Mol Biol 2005, 12(1): 82–89.

54. Barfod, E. T. & Cech, T. R. Deletion of nonconserved helices near the 3’ end of the rRNA intron of Tetrahymena thermophila alters self-splicing but not core catalytic activity. Genes Dev 1988, 2(6): 652–663.

55. Laggerbauer, B., Murphy, F. L. & Cech, T. R. Two major tertiary folding transitions of the Tetrahymena catalytic RNA. The EMBO Journal 1994, 13(11): 2669–2676.

56. Denesyuk, N. A. & Thirumalai, D. How do metal ions direct ribozyme folding? Nat Chem 2015, 7(10): 793–801.

57. Stahley, M. R. & Strobel, S. A. Structural evidence for a two-metal-ion mechanism of group I intron splicing. Science 2005, 309(5740): 1587–1590.

58. Piccirilli, J. A., Vyle, J. S., Caruthers, M. H. & Cech, T. R. Metal ion catalysis in the Tetrahymena ribozyme reaction. Nature 1993, 361(6407): 85–88.

59. Shan, S., Yoshida, A., Sun, S., Piccirilli, J. A. & Herschlag, D. Three metal ions at the active site of the Tetrahymena group I ribozyme. Proc. Natl. Acad. Sci. USA 1999, 96(22): 12299–12304.

60. Yoshida, A., Sun, S. & Piccirilli, J. A. A new metal ion interaction in the Tetrahymena ribozyme reaction revealed by double sulfur substitution. Nat Struct Biol 1999, 6(4): 318–321.

61. Kuo, L. Y. & Piccirilli, J. A. Leaving group stabilization by metal ion coordination and hydrogen bond donation is an evolutionarily conserved feature of group I introns. Biochimica et Biophysica Acta (BBA) - Gene Structure and Expression 2001, 1522(3): 158–166.

62. Weinstein, L. B., Jones, B. C., Cosstick, R. & Cech, T. R. A second catalytic metal ion in group I ribozyme. Nature 1997, 388(6644): 805–808.

63. Lipchock, S. V. & Strobel, S. A. A relaxed active site after exon ligation by the group I intron. Proc. Natl. Acad. Sci. USA 2008, 105(15): 5699–5704.

64. Serganov, A., Huang, L. & Patel, D. J. Coenzyme recognition and gene regulation by a flavin mononucleotide riboswitch. Nature 2009, 458(7235): 233–237.

65. Wilt, H. M., Yu, P., Tan, K., Wang, Y. X. & Stagno, J. R. FMN riboswitch aptamer symmetry facilitates conformational switching through mutually exclusive coaxial stacking configurations. J Struct Biol X 2020, 4: 100035.

66. Schön, P. Imaging and force probing RNA by atomic force microscopy. Methods 2016, 103: 25–33.

67. Chen, Y. & Pollack, L. SAXS studies of RNA: structures, dynamics, and interactions with partners. Wiley Interdiscip Rev RNA 2016, 7(4): 512–526.

68. Lilley, D. M. Analysis of branched nucleic acid structure using comparative gel electrophoresis. Q Rev Biophys 2008, 41(1): 1–39.

69. Liu, Y., Gonen, S., Gonen, T. & Yeates, T. O. Near-atomic cryo-EM imaging of a small protein displayed on a designed scaffolding system. Proc. Natl. Acad. Sci. USA 2018, 115(13): 3362–3367.

70. Liu, Y., Huynh, D. T. & Yeates, T. O. A 3.8 A resolution cryo-EM structure of a small protein bound to an imaging scaffold. Nat Commun 2019, 10(1): 1864.

71. Warner, K. D., Hajdin, C. E. & Weeks, K. M. Principles for targeting RNA with drug-like small molecules. Nat. Rev. Drug Discov. 2018, 17(8): 547–558.

