## Supplementary Information for "Sub-3 Å cryo-EM structure of RNA enabled by engineered homomeric self-assembly"

### Table of Contents

*Materials and Methods*...2
*Supplementary Figure 1*...5
*Supplementary Figure 2*...5
*Supplementary Figure 3*...6
*Supplementary Figure 4*...7
*Supplementary Figure 5*...8
*Supplementary Figure 6*...11
*Supplementary Figure 7*...12
*Supplementary Figure 8*...14
*Supplementary Figure 9*...14
*Supplementary Figure 10*...15
*Supplementary Figure 11*...16
*Supplementary Figure 12*...17
*Supplementary Figure 13*...18
*Supplementary Table 1*...19
*Supplementary Table 2*...20
*Supplementary Table 3*...21
*Supplementary references*...22

### Materials and Methods

**RNA construct design.** The software NanoTiler[1] (developed by Bruce Shapiro's group) was used to design the RNA constructs with kissing-loops (KLs) installed for homomerization. Throughout this work, we chose the 7-bp KL that is adapted from the RNA I-RNA II complex of the *E. coli ColE1* plasmid[2] and has been determined by NMR[3] (PDB code: 2bj2) to have an included angle of ~120°. For TetGI-D and TetGI-DS, the design was based on crystal structure[4] of the core domains of the TetGI (PDB code: 1x8w). Before being imported into NanoTiler as structural modules, the PDB files of 1x8w (the TetGI core) and 2bj2 (the 7-bp KL) were first edited so that the helices (P6b and P8 for the TetGI core and the two stems of the 7-bp KL) to be connected in the final construct were trimmed to 3 bps. Then NanoTiler's command "clone" was used to generate another copy of these structural models; commands "genhelixconstraint" and "opthelices" were used to generate ideal A-form RNA helices of certain lengths for connecting the TetGI core and the 7-bp KL; the "start_score" in the output of the "opthelices" for placing the last connecting helices indicates how good the ring closure is, with a lower value indicating better ring closure. The processes can be automated with the "NanoScript" of NanoTiler in which "foreach" loops can be incorporated to iterate through connecting helices of different lengths to screen for the optimal lengths for ring closure. Similarly, the trimeric construct TetGI-T was designed based on the computational model[5] of the complete TetGI kindly provided by Prof. Eric Westhof; AzoGI-T based on the crystal structure[6] of the *Azoarcus* group I intron (AzoGI) (PDB code: 1u6b); and FMNrsw-T based on the crystal structure[7] of the FMN riboswitch (FMNrsw) bound to FMN (PDB code: 3f2q). After installing the KL sequence and connector helices onto the peripheral stem-loops of each target RNA, the secondary structure of each newly formed stem-loop was further checked by Mfold[8] to ensure that the stem-loop folds correctly.

**RNA preparation and nanostructure assembly.** All RNA molecules were synthesized by *in vitro* transcription (IVT) using the HiScribe™ T7 High Yield RNA Synthesis Kit from the New England Biolabs (NEB). The corresponding DNA templates for *in vitro* transcription were the PCR products of gene fragments (sequences shown in Supplementary table 2) ordered from WuXi Qinglan Biotech. The PCR experiments were conducted using the Q5® Hot Start High-Fidelity DNA Polymerase (NEB) following the recommended protocol provided by NEB. Primers and other modified oligonucleotides were ordered from the Integrated DNA Technologies (IDT). All IVT RNA molecules were purified by denaturing PAGE (containing 7M urea), then ethanol precipitated and suspended in pure water. The RNA concentration was determined by measuring $OD_{260}$.

Protocols for RNA nanostructure assembly were adapted from a previous publication[9]. RNAs were first denatured at 85 °C for 1 min and snap-cooled on ice. Then, the annealing buffers containing 20 mM of Tris-acetate (pH 8.0) and varied concentrations of $Mg^{2+}$ (using 100 mM $Mg^{2+}$ stock solution containing 110 mM of $MgCl_2$ and 10 mM of EDTA) and $Na^+$ (using 1 M $Na^+$ stock solution containing 1 M of NaCl) were added to the denatured RNAs (to a final RNA concentration of ~800 nM). The mixtures were then annealed from 70 °C to 4 °C with the following protocol: 70 °C to 50 °C over 6 min, 50 °C to 37 °C over 20 min, 37 °C to 4 °C over 2 hr. The annealed RNAs were analyzed by native polyacrylamide gel electrophoresis (PAGE) in 0.5 × TBE buffer supplemented by 3 mM of $MgCl_2$ (so that the final concentration of $Mg^{2+}$ is 2 mM because 1 mM of EDTA is included in 0.5 × TBE). To prepare the samples for cryo-EM experiments, the cation contents in the annealing buffers were chosen based on the native PAGE results: for TetGI-DS, TetGI-D, TetGI-T and AzoGI-T, the annealing buffer contains 1 mM of $Mg^{2+}$ and 1.2 equivalents (equiv.) of associated synthetic oligonucleotides (dTetCIRC for TetGI-DS; TetLEM for TetGI-D and -T; and AzoLEM for AzoGI-T); for FMNrsw-T, the annealing buffer contains 0.3 mM of $Mg^{2+}$ and 10 mM of $Na^+$ supplemented by 1 mM of FMN. The assembled nanostructures were purified by preparative native PAGE, eluted to the annealing buffer of 3 mM $Mg^{2+}$, and concentrated to ~1 µg/µL using Amicon Ultra centrifugal filters (MWCO 30 kDa).

**Cryo-EM sample preparation and data acquisition.** Before grid preparation, $Mg^{2+}$ and other components (such as short oligonucleotides or ligand) were added to the purified and concentrated RNA assemblies. For the four ribozyme constructs (TetGI-DS, TetGI-D, TetGI-T and AzoGI-T), $Mg^{2+}$ was added to a final concentration of 30 mM to ensure the complete folding[10]. Additional synthetic oligonucleotide was added for each of the ribozyme construct in case of the dissociation due to equilibrium or during the purification processes: for TetGI-DS, 1.2

equiv. of dTetCIRC was added; for TetGI-D and -T, 3 equiv. of TetLEM was added; and for AzoGI-T, 3 equiv. of AzoLEM was added. For FMNrsw-T, $Mg^{2+}$ was added to a final concentration of 10 mM, and FMN to a final concentration of 200 μM. After the addition of $Mg^{2+}$ and other components, the samples were left at room temperature for 15 min before grid preparation.

Each RNA sample was applied to a glow-discharged Quantifoil (R1.2/1.3, 400 mesh) holey carbon grid. The three TetGI samples were vitrified in liquid ethane with a Thermo Fisher Vitrobot Mark IV (blot for ~7s, force +12) and imaged on a Titan Krios microscope equipped with a Gatan K3 camera. The Krios movie stacks were acquired in counting mode, with a physical pixel size of 0.825 Å, total exposure of 47 e/$Å^2$ and at a defocus ranging from -0.8 to -2.0 μm. The AzoGI-T and FMNrsw-T samples were frozen using a Gatan Cryoplunge 3 (blot ~3s) and imaged on an FEI T30 Polara microscope equipped with a K2 summit detector. The Polara micrographs were acquired in super-resolution mode, with 0.62 Å super-resolution pixel size and 52 e/$Å^2$ exposure, with a defocus ranging from -0.8 to -2.0 μm. SerialEM[11] was used for collection of all datasets. More information about cryo-EM data collection can be found in Supplementary table 3.

**Cryo-EM data processing.** The overall processing pipeline was similar for all the datasets. Briefly, movie stacks were binned 2× if collected in super-resolution mode, then motion-corrected with MotionCor2[12]. The contrast transfer function was subsequently computed with CTFFIND4[13]. Particle picking and 2D particle curation were performed with Simplified Application Managing Utilities of EM Labs (https://liao.hms.harvard.edu/samuel), a set of protocols built on the SPIDER[14] image processing system. About 2,000 particles were manually picked to generate 2D initial models, which were used to auto-pick 10% of the micrographs and generate refined 2D templates. The refined templates were then used to auto-pick from all micrographs. The particles were curated with "samtree2dv3.py", which runs iterative principal component analysis (PCA), k-means clustering and multireference alignment. Selected particles were then imported to RELION 3.0[15], which was used for all subsequent processing steps. Briefly, the particles were downsampled and sorted with 3D classification, before unbinning and 3D refinement with enforced symmetry. As the constructs include monomers of uneven integrity, we used symmetry expansion[16] to address the pseudo-symmetry of the particles and separate the most stable monomers. Monomers were then classified through 3D classification without particle alignment, and selected classes were refined using local angle search refinement. Monomer resolution was calculated with the FSC = 0.143 criterion on half maps from independent halves of the dataset, while local resolution was determined with ResMap[17]. All maps were density modified with "phenix.resolve_cryo_em"[18] that is recently incorporated in PHENIX[19], and no model information was provided to avoid any possible bias. More information about cryo-EM data processing can be found in Supplementary figures 4,6,7,11 and 13.

**Structure modelling.** To build the structural model of the TetGI, we began with the core domains by rigid-body fitting the crystallographic model of the core (PDB code: 1x8w) into the TetGI-DS map using UCSF Chimera[20] and performing several rounds of real-space refinements[21] in PHENIX[19]. Though the crystallographic model mostly agreed with our cryo-EM map, two regions of the core show the most significant discrepancy—the region of P9.0 and G414, which were not present in the crystallographic model, and the region near J8/7, which we believe was built incorrectly in the crystallographic model. These two regions were manually built in COOT[22]. To build the peripheral domains of TetGI, ideal A-form helices were generated and fitted to the TetGI-DS map using UCSF Chimera[20] based on the known secondary structure. The rest of structure was manually built in COOT[22]. The most difficult region to build was P2.1 to P13, where the map density was relatively weaker and the secondary structure assignment was ambiguous. For instance, any residue from A87 through A90 could be possibly unpaired, and the exact number of base-pairs in kissing-loops P13 was not known. We manually tested different possibilities of secondary structure assignments for this region to ensure its successful joining to the helices on both sides. The TetGI-T map was referenced to build the region of P2.1 to P13 because the map quality is better for this region than the TetGI-DS map. The complete TetGI model was finally real-space refined against the maps of the three TetGI constructs and manually checked and adjusted in COOT. The statistics of model refinement and validation were tabulated in Supplementary table 3.

For AzoGI-T, we used the AzoGI crystal structure (PDB code: 1u6b) as the initial model, which was rigid-body fitted to the map to the AzoGI-T map using Chimera, and then the atomic model was iteratively improved by real-space refinements in PHENIX and manual adjustments in COOT. Because the final cryo-EM map for FMNrsw-T is the trimer instead of SE monomer, the designed NanoTiler model was used as the initial model to fit the map by MDFF[23], and then the atomic model was iteratively improved by real-space refinements in PHENIX and manual adjustments in COOT.

**Ribozyme activity assays.** To measure the trans-acting endonucleolytic activity of the TetGI constructs (TetGI-M, TetGI-D and TetGI-T), reactions were performed with 2 µM 56-FAM labelled 10-nt RNA substrate, 0.5 µM ribozyme monomeric units and 1 mM GTP cofactor in 1 × ribozyme reaction buffer (10 mM sodium cacodylate at pH 6.8, 30 mM $MgCl_2$) at 37 °C. For the AzoGI constructs, the activity assayed was based on the second step of splicing because the 3'-exon of the G4-a+7 AzoGI constructs (AzoGI*-M and AzoGI*-T, see Supplementary figure 10) was not completely cleaved during IVT. Reactions were performed with 2 µM 56-FAM labelled 5-nt RNA substrate (mimicking 5'-exon) and 1 µM ribozyme monomeric units in 1 × ribozyme reaction buffer at 25 °C. Aliquots of the reaction mixtures were removed at various times and quenched by adding an equal volume of 2 × TBE–urea sample buffer (Bio-Rad) supplemented with extra 50 mM EDTA. Reactions were analyzed by dPAGE and the gels were visualized using a Sapphire Biomolecular Imager (Azure Biosystems).

**Reverse transcription and polyadenylation assay.** Reverse transcription (RT) was used to study the 5' sequence of the TetGI constructs. The reactions were performed with 0.8 µM RNA, 1.2 µM 56-FAM labelled RT primer (complementary to nucleotides 29 to 46 of the TetGI), 0.5 mM of each dNTP and 1 U/µL AMV reverse transcriptase (NEB) in 1 × AMV Reverse Transcriptase Reaction Buffer (supplied by manufacturer with the enzyme) for 1 hour at 42 °C. The TetGI 5'-mimic RNA (Supplementary figure 2) was prepared by IVT and analyzed by dideoxynucleotide sequencing to provide length markers for determining the 5' cleavage site. For these sequencing reactions, 1.25 mM of a single ddNTP (dideoxynucleoside triphosphate; GE Healthcare) was added to each RT reaction for the 5'-mimic RNA. Polyadenylation assay[24] was used to determine if the RNA has a 3'-hydroxyl group or a 2′-3′ cyclic phosphate group because the former can be polyadenylated while the latter cannot. Polyadenylation reactions were performed with 200 nM RNA, 0.5 mM ATP and 24 U/µL yeast poly(A) polymerase (Thermo Scientific™) in 1 × Poly(A) Polymerase Reaction Buffer (supplied by manufacturer with the enzyme) for 20 min at 37 °C.

**Ligand binding assays for FMNrsw assemblies.** The binding affinities of the FMNrsw assemblies to FMN ligand was measured based on the fluorescence quenching of FMN after specific binding to the riboswitch[7]. Varying concentrations (ranging from 0.1 nM to 100 nM of monomeric units) of the FMNrsw assemblies (monomer control, FMNrsw-T dimer and trimer) were mixed with 60 nM FMN in 50 mM Tris-HCl (pH 7.4), 100 mM KCl and 2 mM $MgCl_2$ for at least 30 min at room temperature for equilibrium before data collection. The fluorescence intensity was measured at 530 nm emission with 450 nm excitation at room temperature using a Synergy H1 Hybrid multi-mode microplate reader (BioTek). Each data point was measured from three independent experiments. Data were fitted using a two-parameter ($K_d$ and $f_c$) quadratic Equation (4) derived in Supplementary figure 12 implying 1:1 stoichiometry of ligand to monomeric unit. Data plotting and curve fitting were performed using OriginPro 2018 software.

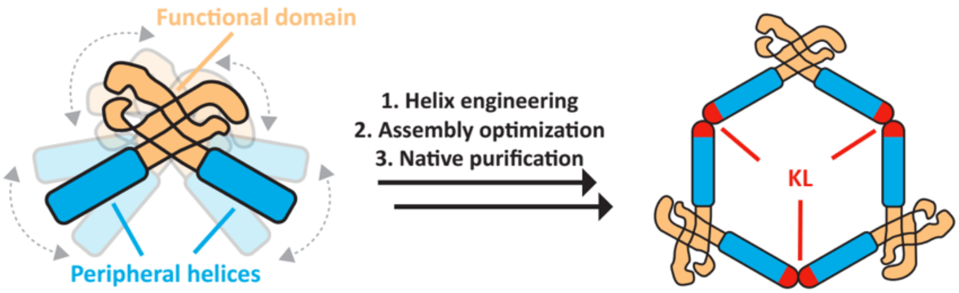


| Properties | Standalone RNA | Assembled RNA |
|---|---|---|
| Molecular weight | Small | Multiplied |
| Conformation | Highly flexible | Flexibility mitigated |
| Folding | Strong misfolding tendency, difficult for purification | Alternate folding eliminated by purification |
| Shape | Irregular | Symmetric, characteristic |

**Supplementary Figure 1 | The concept of RNA oligomerization-enabled cryo-EM via installing kissing-loops (ROCK).** The engineering of a target RNA for the self-assembly of a closed homomeric ring, which, compared to the standalone monomer RNA, is more amenable for cryo-EM structural determination due to the tabulated properties.

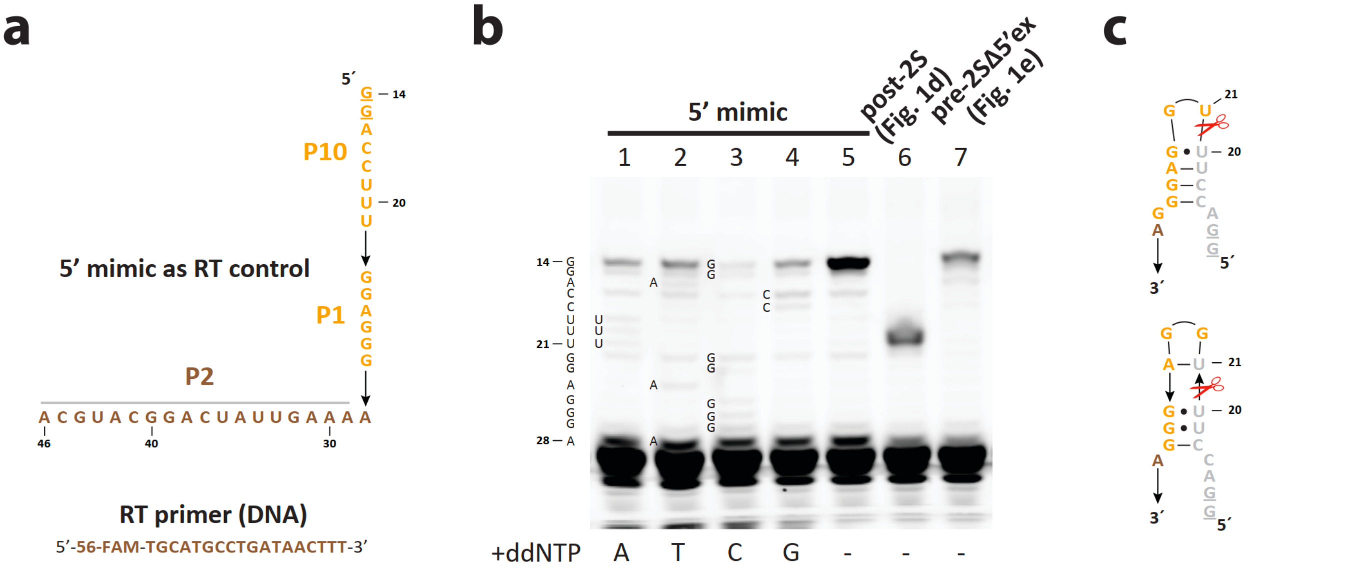


**Supplementary Figure 2 | Self-cleavage of 5'-end of the wild-type TetGI. a**, Sequences of the RNA mimicking the 5'-end of the TetGI (5'-mimic) as the reverse transcription (RT) control and the 56-FAM-labelled DNA primer for RT. In the 5'-mimic, underlined nucleotides were mutated from the wild-type to facilitate the synthesis by IVT; the nucleotides marked by grey line above are the primer-binding region. **b**, Analysis of the 5'-end of the TetGI by RT. Lanes 1 to 5 are the dideoxynucleotide sequencing results of the 5'-mimic with a single dideoxynucleoside triphosphate (ddNTP; indicated under the gel) added (lanes 1 to 4) or no ddNTP added (lane 5); lanes 6 and 7 are the results of the TetGI with wild-type 5' sequence (same as the post-2S complex in Fig. 1d) or mutated 5' sequence (same as the pre-2SΔ5'ex complex in Fig. 1e). While the RNA used in the post-2S complex is cleaved after U20, the RNA in the pre-2SΔ5'ex complex is not cleaved at this site. **c**, The likely formation of hairpin structures (two possibilities shown) of the wild-type 5' sequence accounting for the self-cleavage at the 5'-end. We note that the reaction of the wild-type TetGI leads to a L-19 product that is cleaved after U19[25]. We attribute the difference of cleavage products to the different 5' sequences.

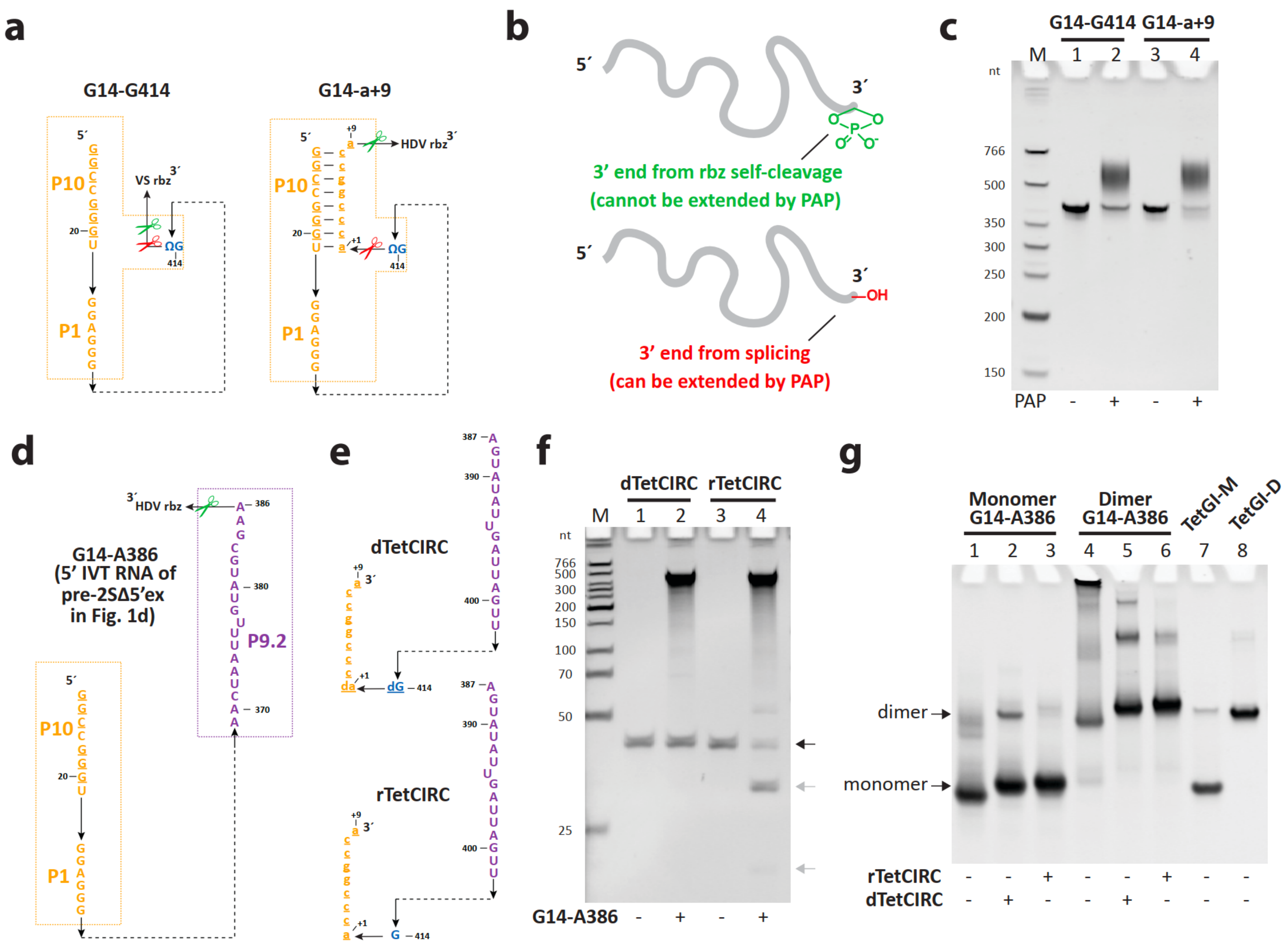


**Supplementary Figure 3 | Designing the 3'-end for the pre-2SΔ5'ex TetGI complex. a**, Two RNA constructs transcribed with a self-cleaving ribozyme (rbz; VS or HDV) for producing a homogeneous 3'-end. The G14-a+9 RNA was originally designed to have 3'-exon (also see G4-a+7 RNA of the AzoGI in Supplementary figure 10), but the 3'-exon was almost completely cleaved in the absence of the 5'-exon. Green and red scissors mark the cleavage sites for the appended ribozyme and the group I intron itself, respectively. **b**, Chemical structures of the 3'-end produced by the cleavage of the appended ribozyme or the splicing of the intron. While the latter can be extended by yeast poly(A) polymerase (PAP), the former cannot. **c**, PAP extension assay demonstrates that the majority products of both G14-A386 (lanes 1 and 2) and G14-a+9 (lanes 3 and 4) RNA preparation are extended by PAP (lanes 2 and 4), indicating that they are mostly the products from the intron splicing or hydrolysis reaction. **d, e**, Sequences of the 5' IVT RNA (**d**; G14-A386, from nucleotides 14 to 386) and the 3' fragments (**e**; dTetCIRC with two deoxynucleotides flanking the 3' splice site, and rTetCIRC with all-RNA nucleotides) for the pre-2SΔ5'ex TetGI complex. **f**, TetGI-catalyzed hydrolysis at the 3' splice site is inhibited by the modification of DNA bases flanking the scissile phosphate of the 3' splice site. Black arrow points to the band of the 3' fragments of (dTetCIRC or rTetCIRC); grey arrows point to the bands of the cleaved 3' fragments. **g**, Assembly assays for the monomeric (lanes 1 to 3) and dimeric (lanes 4 to 6) constructs of 5' IVT G14-A386 RNA without 3' fragment (lanes 1 and 4), with dTetCIRC (lanes 2 and 5) or with rTetCIRC (lanes 3 and 6). The annealing buffer contains 3 mM $Mg^{2+}$, which is chosen from the assembly assay of the post-2S constructs (TetGI-M, -D, and -T; see Fig. 3a). Lanes 7 and 8 are control assemblies of TetGI-M and -D. Without 3' fragment (lanes 1 and 4), there are trailing smears for the 5' IVT RNA, indicating incorrect folding of the intron if P9.2, P9a and P9.0 are not formed. In the presence of dTetCIRC (lanes 2 and 5) or rTetCIRC (lanes 3 and 7), especially for dTetCIRC, some slower-migrating bands emerged, probably due to domain swapping, i.e. a 3' fragment simultaneously binds to a 5' IVT RNA molecule to form P9.2 and another 5' IVT RNA molecule to form P10.

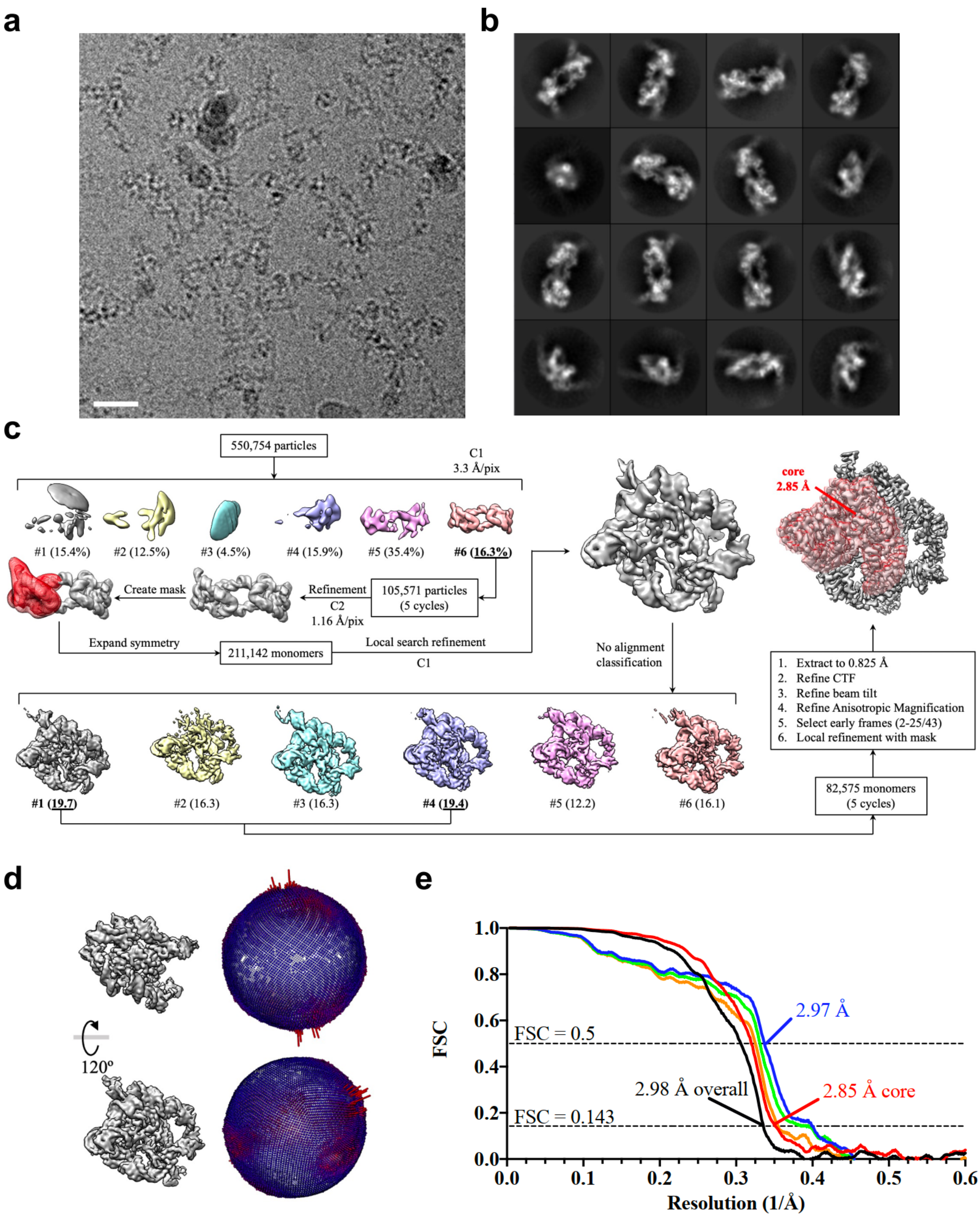


**Supplementary Figure 4 | Cryo-EM imaging, processing and validation for TetGI-DS. a,** Representative cryo-EM image of TetGI-DS. The scale bar represents 20 nm. **b,** 2D class averages of TetGI-DS. Box size is 264 Å. **c,** Processing flowchart for the TetGI-DS dataset. **d,** Angle distribution for the particles included in the final 3D reconstruction. **e,** Fourier Shell Correlation (FSC) curves of the final TetGI-DS reconstruction. Half map #1 vs. half map #2 for the entire monomer is shown in black. The remaining FSC curves were calculated for the core domains only: half map #1 vs. half map #2 (red), model vs. refined map (blue), model refined in half map #1 vs. half map #1 (green), and model refined in half map #1 vs. half map #2 (orange).

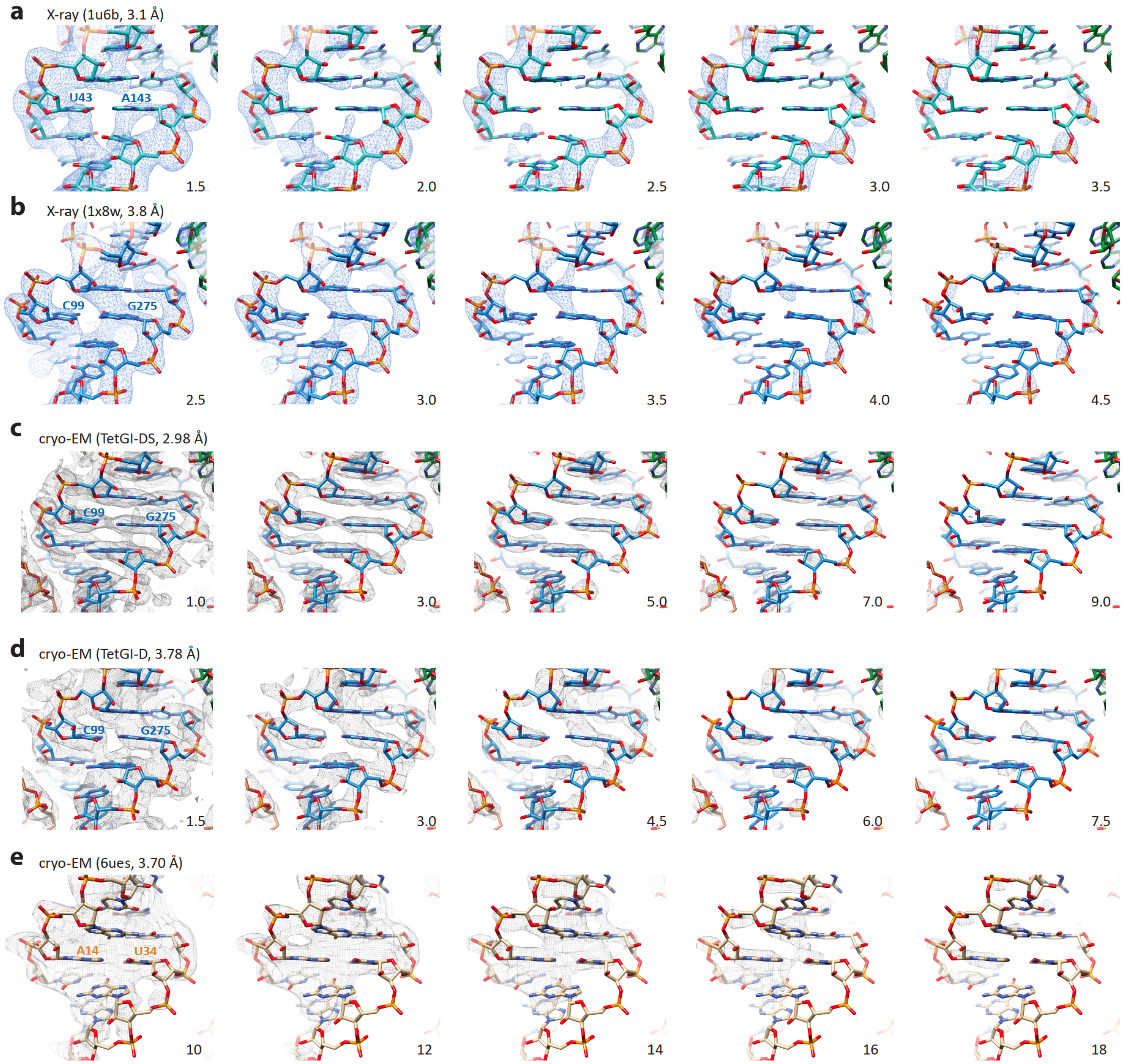


**Supplementary Figure 5 | Comparing features of X-ray and cryo-EM maps at different resolutions. a, b**, Structural models and X-ray maps (blue meshes) of P3 helices of the AzoGI (**a**) and the TetGI (**b**). The maps are generated from structure factors deposited at the PDB (PDB codes and resolutions are given in parentheses). **c, d**, Structural models and EM maps (grey meshes) of P3 helices of TetGI-DS (**c**) and TetGI-D (**d**). **e**, Structural model and EM map (grey meshes) of the P2a helix of the SAM-IV riboswitch in the apo state[26] (PDB code: 6ues) are shown for comparison. The contour levels ($\sigma$ for X-ray maps and RMSD for EM maps) are indicated at the bottom right corner. Unlike the X-ray maps, phosphate groups are not necessarily the most prominent intensity for medium- or low-resolution cryo-EM maps of RNA due to resolution-dependent signal loss for phosphate groups in cryo-EM maps[27]. This underscores the difficulty for interpreting medium- or low-resolution cryo-EM maps of RNA. It is entirely imaginable how difficult it could be to model the regions of an even lower local resolution and/or corresponding to nonhelical structures.

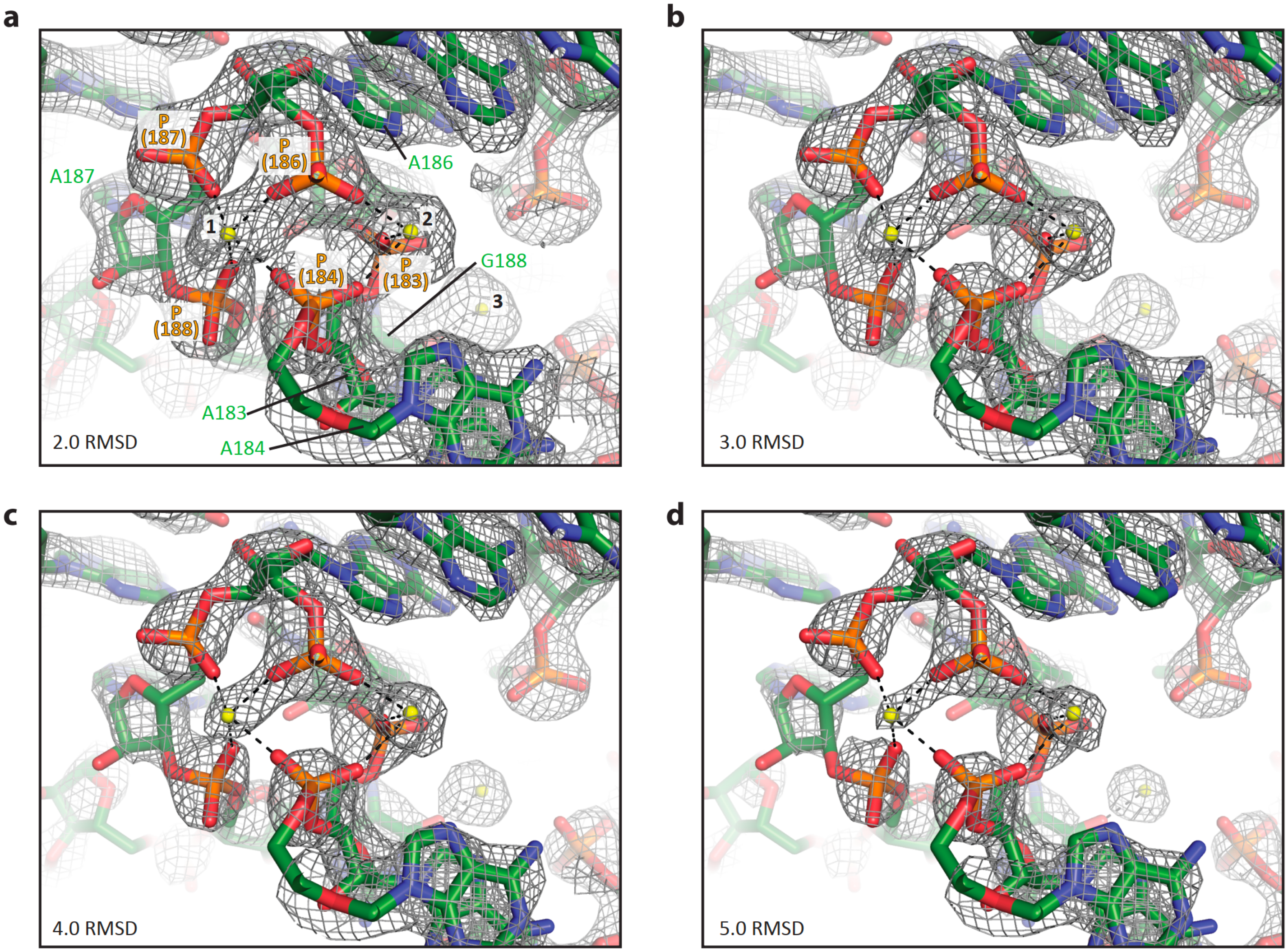


**Supplementary Figure 6 | Three $Mg^{2+}$ ions at the A-rich bulge of P4-P6 domain. a** to **d**, The cryo-EM map of TetGI-DS near the A-rich bulge are shown with different contour levels (indicated at the bottom-left corner of each panel). Consistent with previous studies[28-30], all the three $Mg^{2+}$ (yellow spheres) in this region are observed in our map: #1, coordinating to the phosphate oxygens of A184, A186, A187 and G188; #2, coordinating to the phosphate oxygens of A183, A184 and A186; #3, coordinating to O6 of G188. We note that the map intensity of a $Mg^{2+}$ ion, among other variable factors, is dependent on the number of inner-sphere coordination interactions it forms with RNA; and a higher intensity is observed for a $Mg^{2+}$ ion of a smaller number of inner-sphere coordination interactions with RNA. We attribute this to the fact that the coordinated water molecules contribute to the map intensity because the water molecules cannot be resolved from the $Mg^{2+}$ ions at this resolution. As is shown here, the intensity of #3 is stronger than that of #2, and the intensity of #2 is stronger than that of #1.

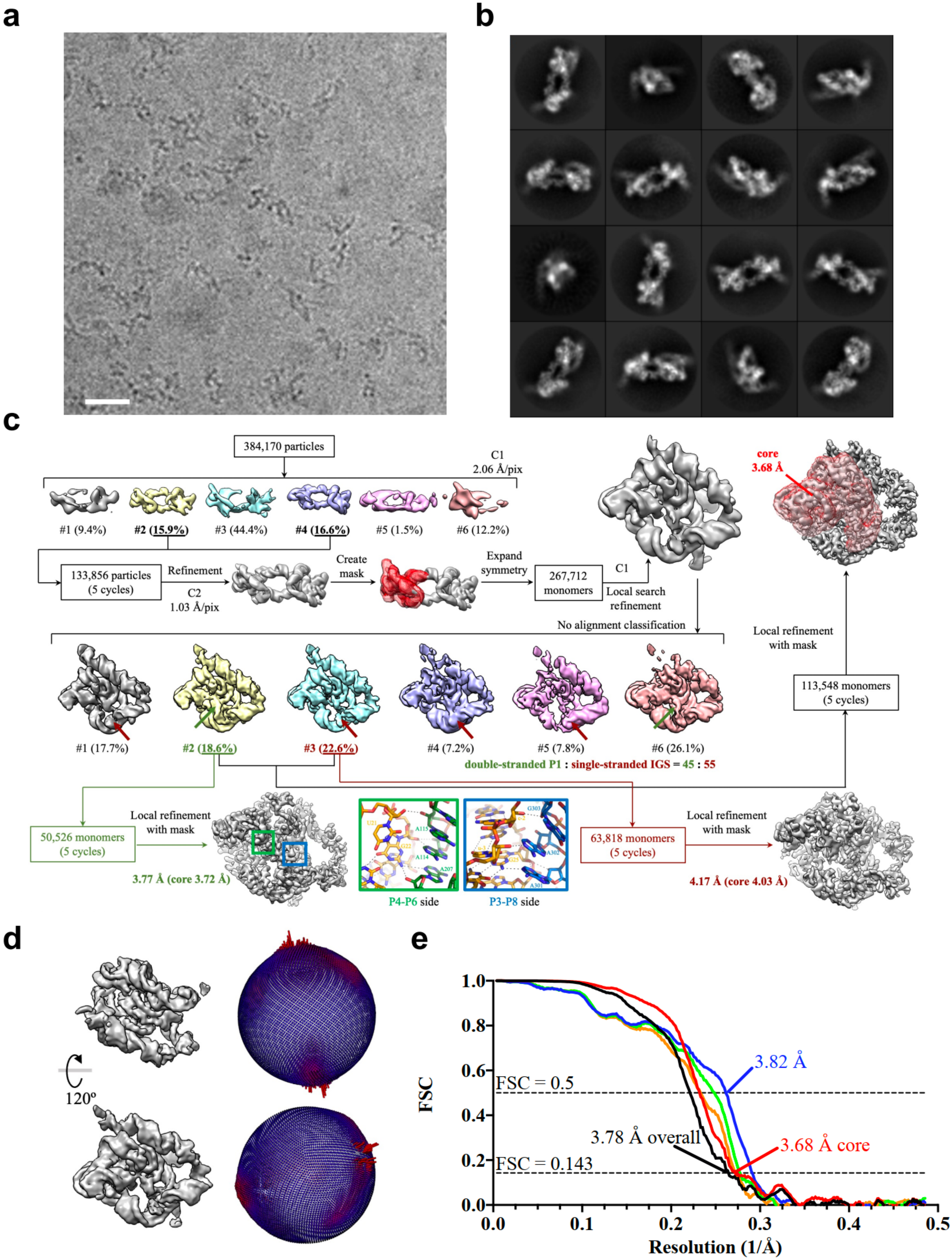

a
b
c
384,170 particles
C1
2.06 Å/pix
#1 (9.4%)
#2 (15.9%)
#3 (44.4%)
#4 (16.6%)
#5 (1.5%)
#6 (12.2%)
133,856 particles
(5 cycles)
Refinement
C2
1.03 Å/pix
Create mask
Expand symmetry
267,712 monomers
C1
Local search refinement
core
3.68 Å
No alignment classification
Local refinement with mask
113,548 monomers
(5 cycles)
#1 (17.7%)
#2 (18.6%)
#3 (22.6%)
#4 (7.2%)
#5 (7.8%)
#6 (26.1%)
double-stranded P1 : single-stranded IGS = 45 : 55
Local refinement with mask
50,526 monomers
(5 cycles)
3.77 Å (core 3.72 Å)
P4-P6 side
P3-P8 side
63,818 monomers
(5 cycles)
Local refinement with mask
4.17 Å (core 4.03 Å)
d
120º
e
FSC
FSC = 0.5
FSC = 0.143
3.82 Å
3.78 Å overall
3.68 Å core
Resolution (1/Å)

**Supplementary Figure 7 | Cryo-EM imaging, processing and validation for TetGI-D. a,** Representative cryo-EM image of TetGI-D. The scale bar represents 20 nm. **b,** 2D class averages of TetGI-D. Box size is 264 Å. **c,** Processing flowchart for the TetGI-D dataset. 3D classification of the symmetry-expanded monomers results in classes according to the conformations with double-stranded P1 (green arrows) or with single-stranded IGS (red arrows). The ratio of the two conformations is calculated and shown. For the conformation with double-stranded P1, green and blue boxes show the details of tertiary contacts of P1 on the P4-P6 side and P3-P8 side, respectively. The contacts agree well with the previous biochemical studies[31-33] and the crystal structure of AzoGI[6]. The final cryo-EM map for TetGI-D was refined from two classes, and exhibits a stronger map intensity of double-stranded P1 than single-stranded IGS; therefore, the atomic model was built with double-stranded P1. **d,** Angle distribution for the particles included in the final 3D reconstruction. **e,** Fourier Shell Correlation (FSC) curves of the final TetGI-D reconstruction. Half map #1 vs. half map #2 for the entire monomer is shown in black. The remaining FSC curves were calculated for the core domains only: half map #1 vs. half map #2 (red), model vs. refined map (blue), model refined in half map #1 vs. half map #1 (green), and model refined in half map #1 vs. half map #2 (orange).

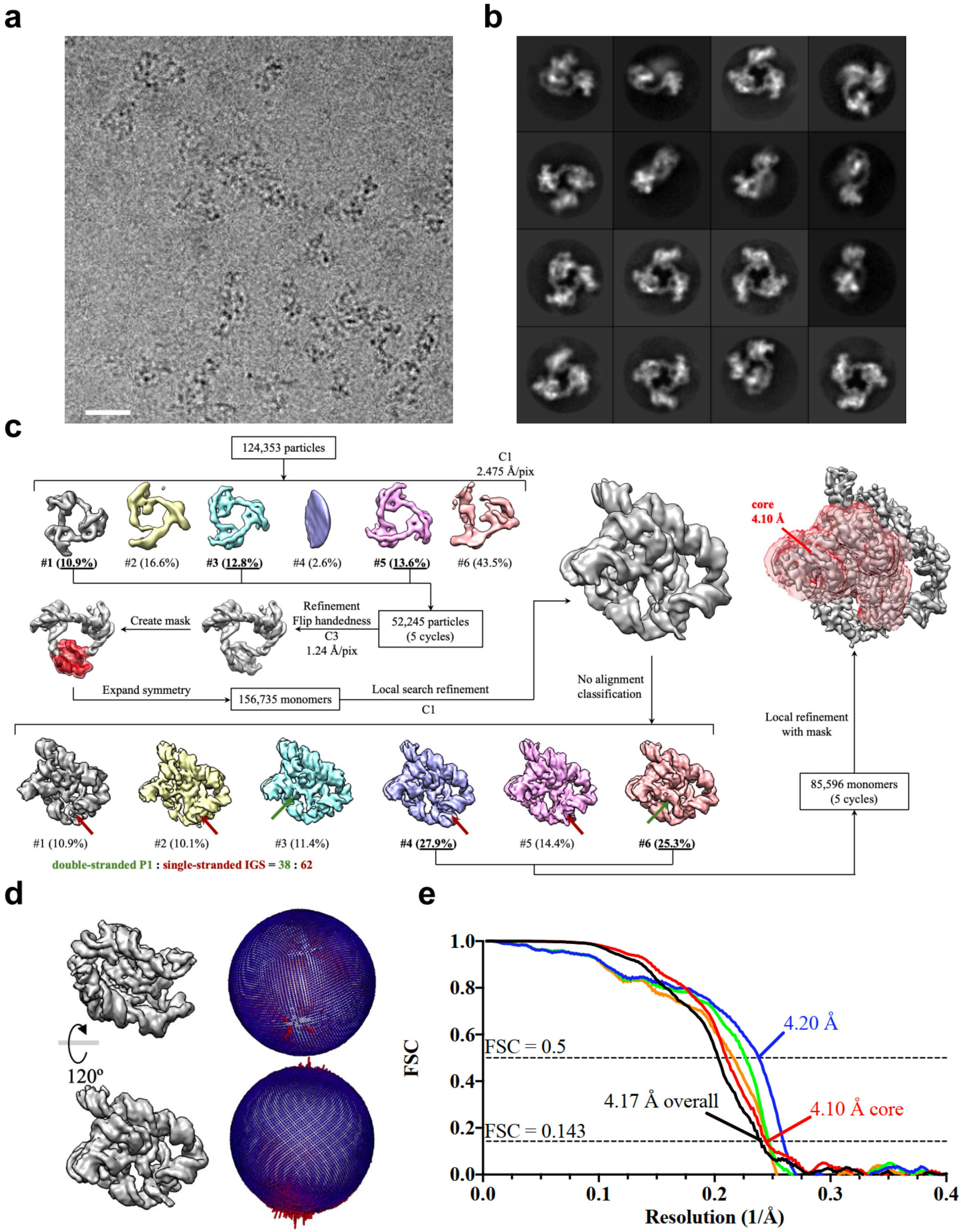


**Supplementary Figure 8 | Cryo-EM imaging, processing and validation for TetGI-T. a,** Representative cryo-EM image of TetGI-T.

The scale bar represents 20 nm. **b,** 2D class averages of TetGI-T. Box size is 317 Å. **c,** Processing flowchart for the TetGI-T dataset. Similar to the case of TetGI-D, 3D classification of the symmetry-expanded monomers of TetGI-T also results in classes according to the conformations with double-stranded P1 (green arrows) or with single-stranded IGS (red arrows). The ratio of the two conformations is calculated and shown, which is close to the dimeric construct TetGI-D shown in Supplementary figure 6. The final cryo-EM map for TetGI-T was refined from two classes, and exhibits a stronger map of single-stranded IGS than double-stranded P1, probably due to the stabilization of the peripheral domains by the trimer construct; therefore, the atomic model was built with the single-stranded IGS. **d,** Angle distribution for the particles included in the final 3D reconstruction. **e,** Fourier Shell Correlation (FSC) curves of the final TetGI-T reconstruction. Half map #1 vs. half map #2 for the entire monomer is shown in black. The remaining FSC curves were calculated for the core domains only: half map #1 vs. half map #2 (red), model vs. refined map (blue), model refined in half map #1 vs. half map #1 (green), and model refined in half map #1 vs. half map #2 (orange).

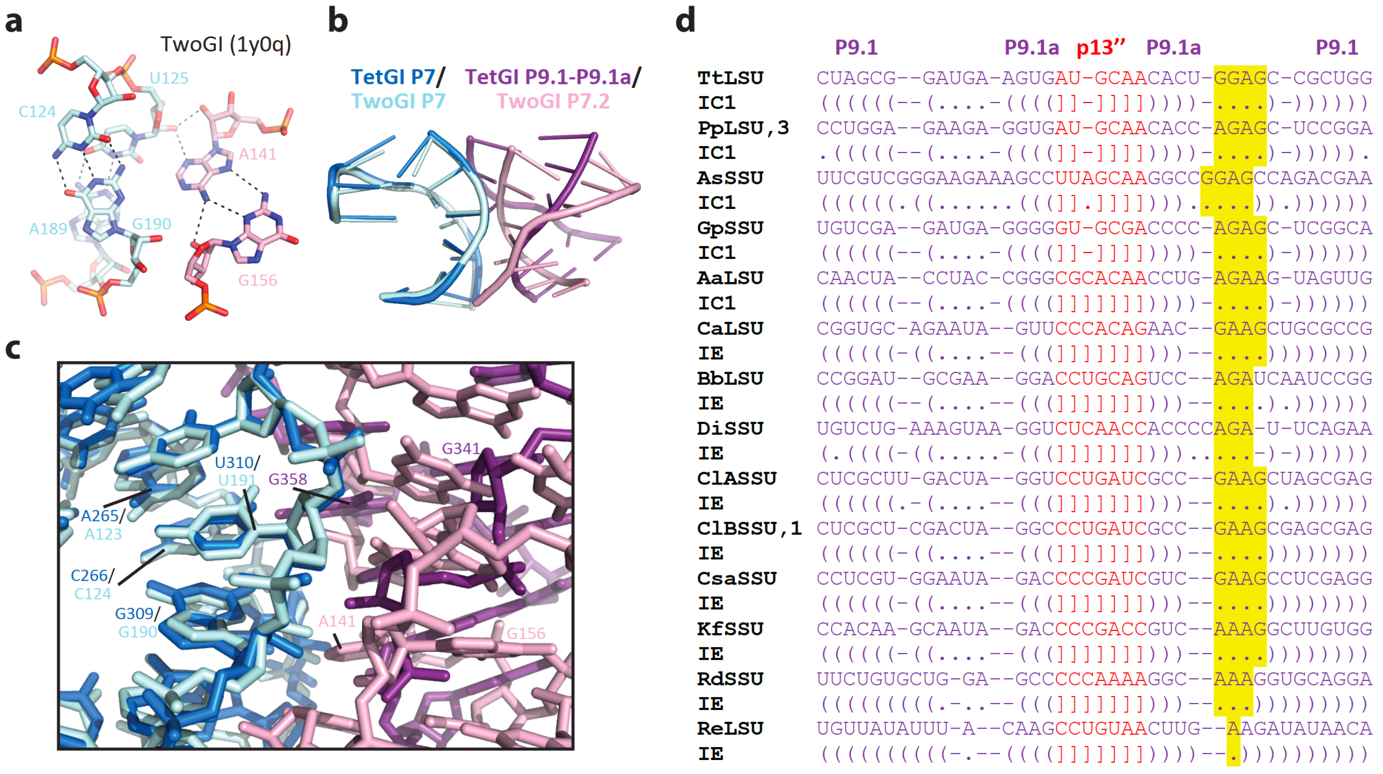


**Supplementary Figure 9 | A conserved long-range tertiary interaction involving P7 and the peripheral domain.** **a**, A similar contact is observed in the crystal structure of the Twort group I intron[34] (TwoGI; PDB code: 1y0q) formed between P7 and the internal loop in P7.2. **b**, **c**, Comparing the tertiary interactions observed in TetGI and TwoGI (overlayed by P7). **d**, Sequence alignment of different class IC1 and class IE group I introns reveals a conserved purine-rich loop (highlighted in yellow) at J9.1/9.1a. The extracts of the alignments of 14 sequences are from Lehnert et al.[5], where all the sequences were regarded as subgroup IC1, but later some of them were categorized as subgroup IE[35]. TtLSU (intron in the large ribosomal RNA precursor of *Tetrahymena thermophila*) is the TetGI studied in this work. We note that the internal loop of J9.1/9.1a of the TetGI, which has the sequence of 5’-AUGA-3’/5’-GGAG-3’, is reminiscent of but not exactly the loop E motif[36], which is normally 5’-AGUA-3’/5’-GAA-3’, though the sequence of the abovementioned internal loop in P7.2 of the TwoGI is the same as loop E motif.

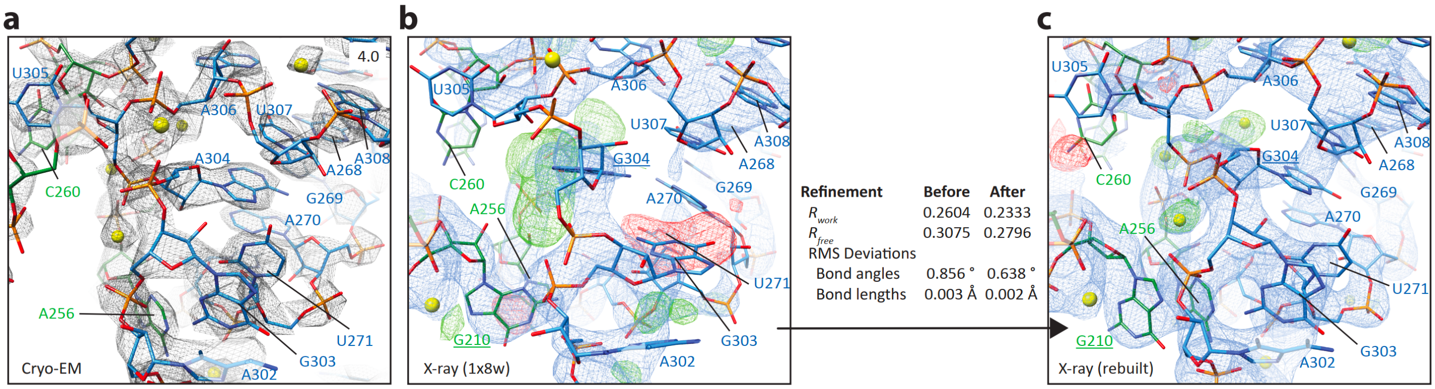


**Supplementary Figure 10 | Rebuilding the crystal structure of the TetGI core using the cryo-EM model.** The high-resolution cryo-EM structure of TetGI-DS (**a**) of the area around J8/7 enables the rebuilding and re-refinement of the corresponding area of the previous crystal structure of the TetGI core[4] (**b** and **c** show the maps and models before and after rebuilding, respectively; $2F_o$−$F_c$ map shown in blue mesh contoured at 2.5σ level, and $F_o$−$F_c$ map shown in green/red meshes contoured at 4σ level). Underlined nucleotides indicate the mutations introduced into the construct for crystallographic study. The crystallographic maps were generated using the factors deposited with the PDB (code: 1x8w). There are four RNA molecules in the asymmetric unit of the 1x8w crystal and the shown region is from molecule B. To eliminate the potential bias of using a different refinement software, the initial maps and model before rebuilding were also re-refined using phenix.refine and the statistics also improved compared to the data originally deposited with the PDB. The improved refinement statistics may also be attributed to improvement through the RNA chains besides J8/7 region.

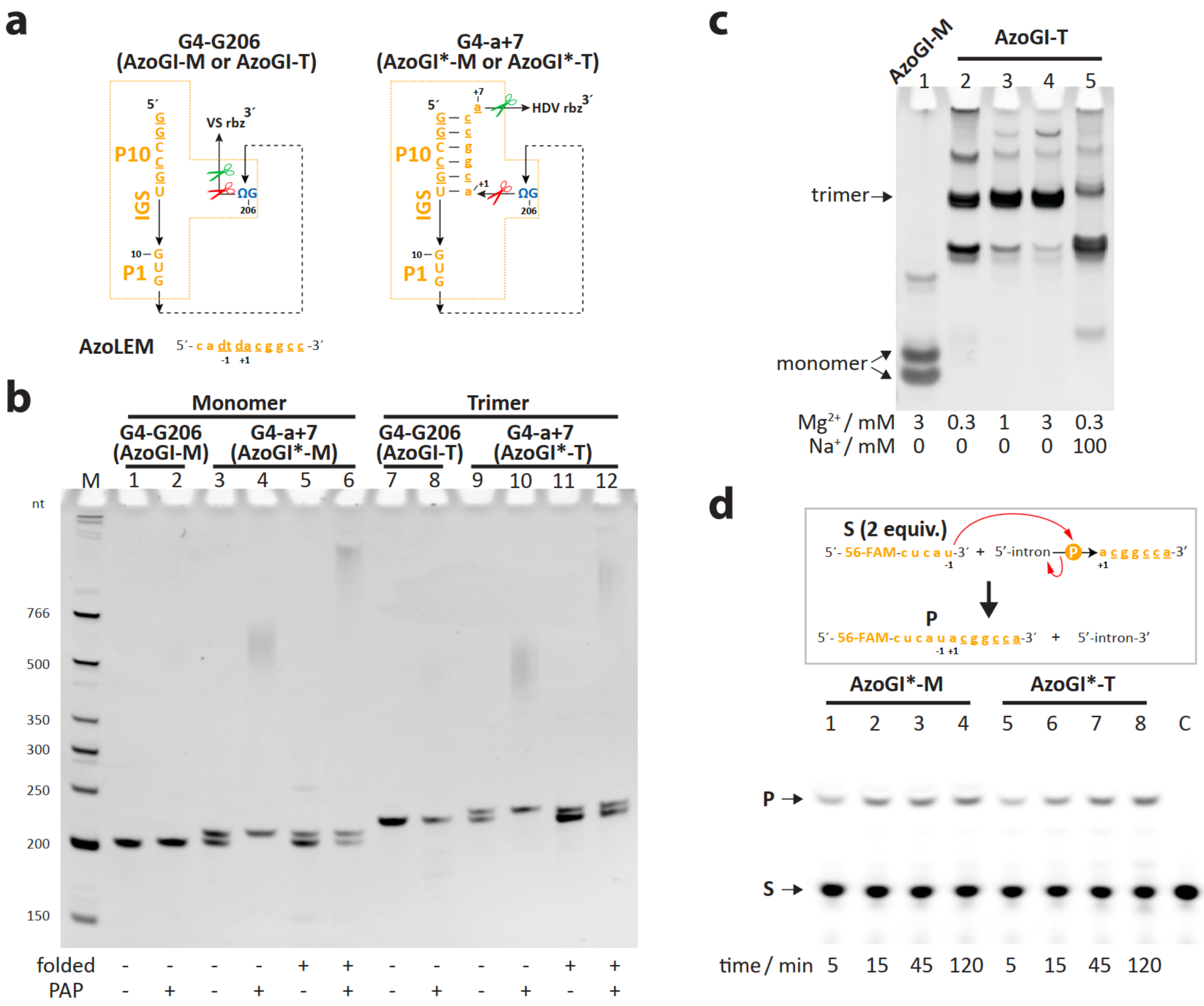


**Supplementary Figure 11 | AzoGI-T sequence design, assembly and activity. a**, Two RNA designs, G4-G206 and G4-a+7, transcribed with a self-cleaving ribozyme (rbz) for producing a homogeneous 3' end. Green and red scissors mark the cleavage site for the appended ribozyme or the intron itself. AzoLEM is the ligated exon mimic for the AzoGI, and its complexing with G4-G206 RNA forms the post-2S state of the intron. **b**, The PAP extension assay of various constructs for the AzoGI. The products from the preparation of G4-G206 RNA constructs cannot be extended by PAP (lanes 2 and 8), indicting a 2',3'-cyclic phosphate generated by rbz at the 3' end (this is different from the G14-G414 RNA of TetGI, the majority of which can be extended by PAP; see lane 2 of Supplementary figure 3c). For G4-a+7 constructs, the products generated by the cleavage of rbz and intron itself can be directly distinguished based on the different electrophoretic mobilities in the presented analytical gel because the former is longer due to the appendage of a 7-nt 3' exon (the small size difference is not noticeable in the preparative gel for RNA purification). As shown in lanes 3 and 9, about half of G4-a+7 RNA of either construct is cleaved at the 3' splice site (whereas, in the case of the TetGI G14-a+9 RNA, more than 90% is cleaved at the 3' splice site, indicating a substantially higher activity of the TetGI; see lane 4 of Supplementary figure 3c). Only the shorter products, which is produced by intron cleavage, can be extended by PAP. After folding, the intron-cleaved shorter products increase to ~70% (lanes 5 and 11), and some portion of these shorter products cannot be extended by PAP, probably due to the lower accessibility of the 3' end after RNA folding. **c**, Assembly assay of the trimeric construct AzoGI-T (lanes 2 to 5) and the monomer control AzoGI-M (lane 1). Similar to the TetGI constructs, the optimal condition for folding/assembly was determined to be 3 mM $Mg^{2+}$. Interestingly, monomer control AzoGI-M runs into two major bands, indicating conformational heterogeneity. **d**, Activity assay to test the activity of the second step of splicing. The reactions were conducted with 1 μM monomer units of either intron construct and 2 μM substrate (S, reacting as the 5' exon) at 25 °C. The assay takes the advantage of the fact that there is still about 30% of AzoGI*-M and AzoGI*-T constructs containing 5' exon after IVT preparation, purification and folding due to presumably lower splicing activity of the AzoGI.

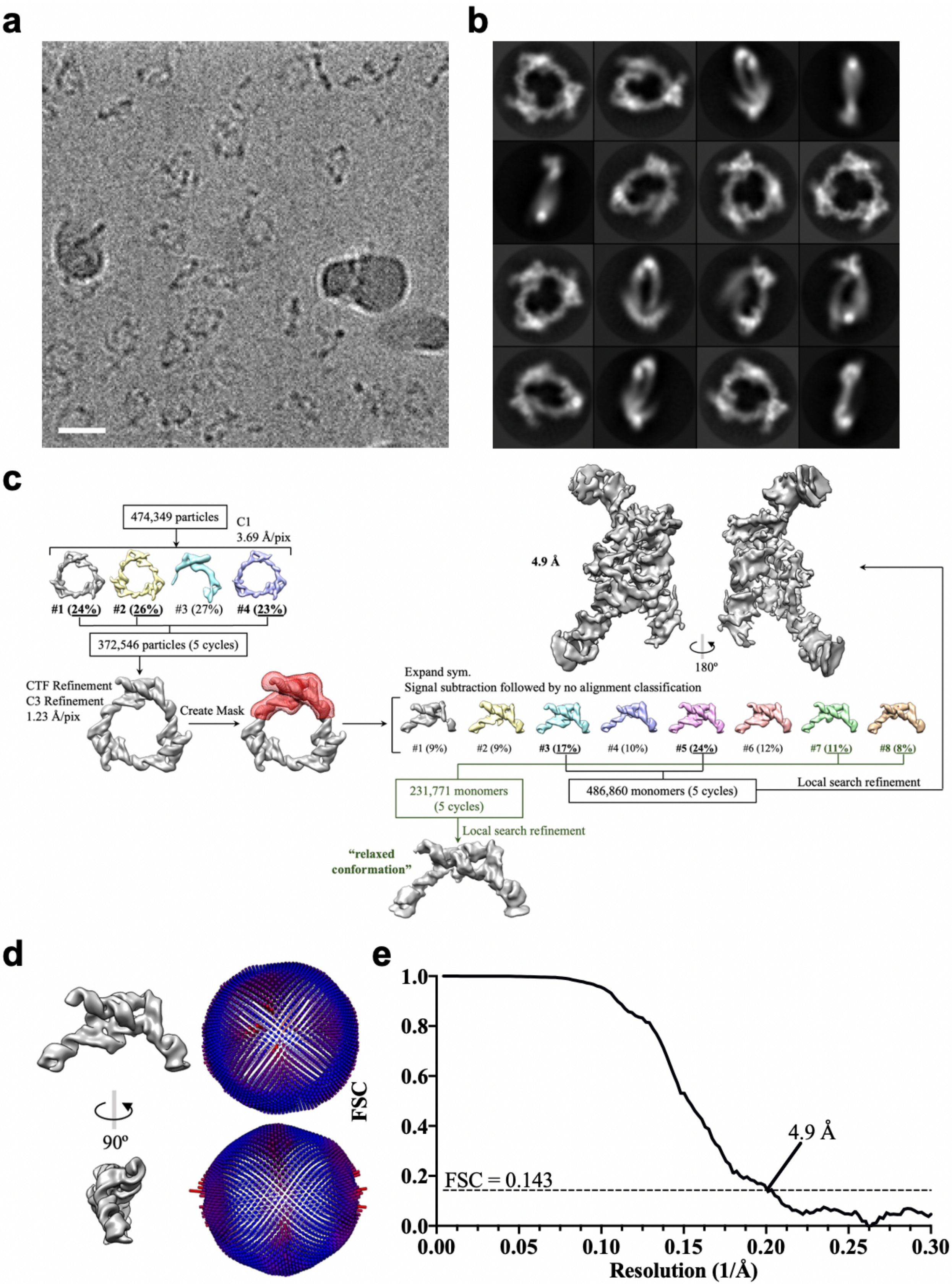


**Supplementary Figure 12 | Cryo-EM imaging, processing and validation for AzoGI-T. a,** Representative cryo-EM image of AzoGI-T. The scale bar represents 20 nm. **b,** 2D class averages of AzoGI-T. Box size is 236 Å. **c,** Processing flowchart for the AzoGI-T dataset. **d,** Angle distribution for the particles included in the final 3D reconstruction. **e,** Gold-standard Fourier Shell Correlation (FSC) for the final AzoGI-T reconstruction.

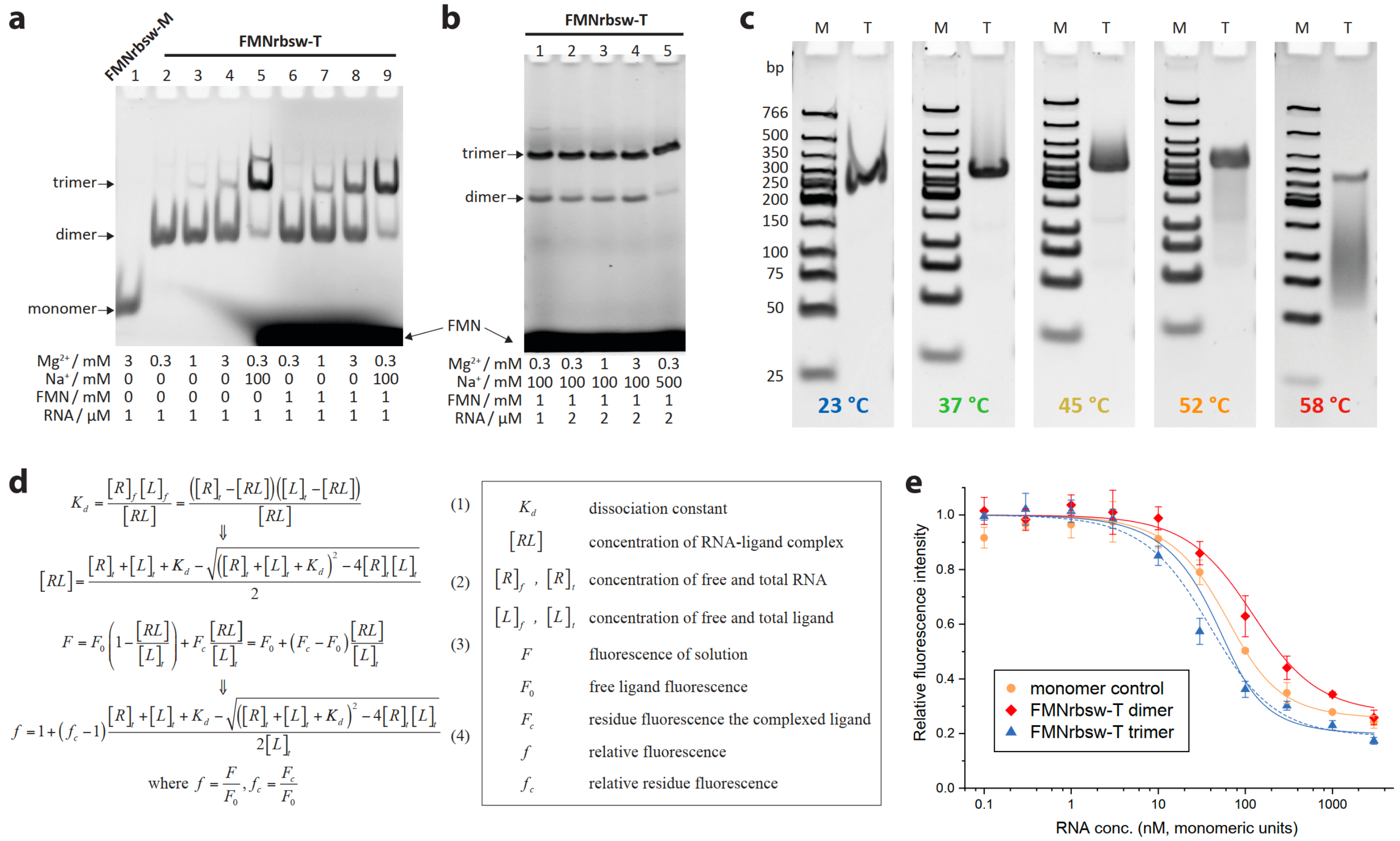


$$K_d = \frac{[R]_f[L]_f}{[RL]} = \frac{([R]_t-[RL])([L]_t-[RL])}{[RL]} \quad (1)$$

$$\Downarrow$$

$$[RL] = \frac{[R]_t+[L]_t+K_d-\sqrt{([R]_t+[L]_t+K_d)^2-4[R]_t[L]_t}}{2} \quad (2)$$

$$F = F_0\left(1-\frac{[RL]}{[L]_t}\right)+F_c\frac{[RL]}{[L]_t} = F_0+(F_c-F_0)\frac{[RL]}{[L]_t} \quad (3)$$

$$\Downarrow$$

$$f = 1+(f_c-1)\frac{[R]_t+[L]_t+K_d-\sqrt{([R]_t+[L]_t+K_d)^2-4[R]_t[L]_t}}{2[L]_t} \quad (4)$$

$$\text{where } f=\frac{F}{F_0},\ f_c=\frac{F_c}{F_0}$$

**Supplementary Figure 13 | FMNrsw-T assembly and activity. a**, **b**, Assembly assay. FMNrsw-M is the monomer control. Three factors to increase the trimer yield: the presence of FMN ligand; high $Na^+$; and increasing RNA concentration. **c**, Native PAGE (6%, in the presence of 2 mM free $Mg^{2+}$) analysis of preassembled FMNrsw-T trimer (lane T) under different temperatures ranging from 23°C to 58°C. Disassembly of the trimer starts to occur at 45 °C. The high thermal stability of the KL may contribute to the formation of the kinetic assembly of dimer because the KL formation may precede the complete folding of the riboswitch during annealing as suggested by its weaker ligand binding as shown in **e**. Lane M contains the low-molecular weight DNA ladder (NEB). **d**, Derivation of the equation to analyze the ligand binding (1:1 stoichiometry) based on fluorescence quenching. Data were fitted using a two-parameter ($K_d$ and $f_c$) quadratic Equation (4). **e**, Fluorescent binding assay of FMN (60 nM) with the riboswitch conducted in 100 mM KCl and 2 mM $MgCl_2$. FMNrbsw-T dimer is the dimeric assembly of the FMNrbsw-T RNA. The calculated $K_d$ (nM, mean ± s.d.), $f_c$ (mean ± s.d.), and $R^2$ are: monomer control, 30.1 ± 2.2, 0.254 ± 0.006, 0.9961; FMNrbsw-T dimer, 91.6 ± 21.9, 0.273 ± 0.017, 0.9944; FMNrbsw-T trimer, 17.4 ± 7.6, 0.197 ± 0.020, 0.9881. The dashed blue line for is fitted FMNrbsw-T trimer taking $[L]_t$ as 20 nM, which assumes that each monomeric subunit in the trimer can locally sense only one third of the ligand, and calculated values of $K_d$ (nM, mean ± s.d.), $f_c$ (mean ± s.d.), and $R^2$ are: 28.8 ± 6.3, 0.188 ± 0.015, 0.9937. This indeed results in an improved fitting as reflected by an improved $R^2$.

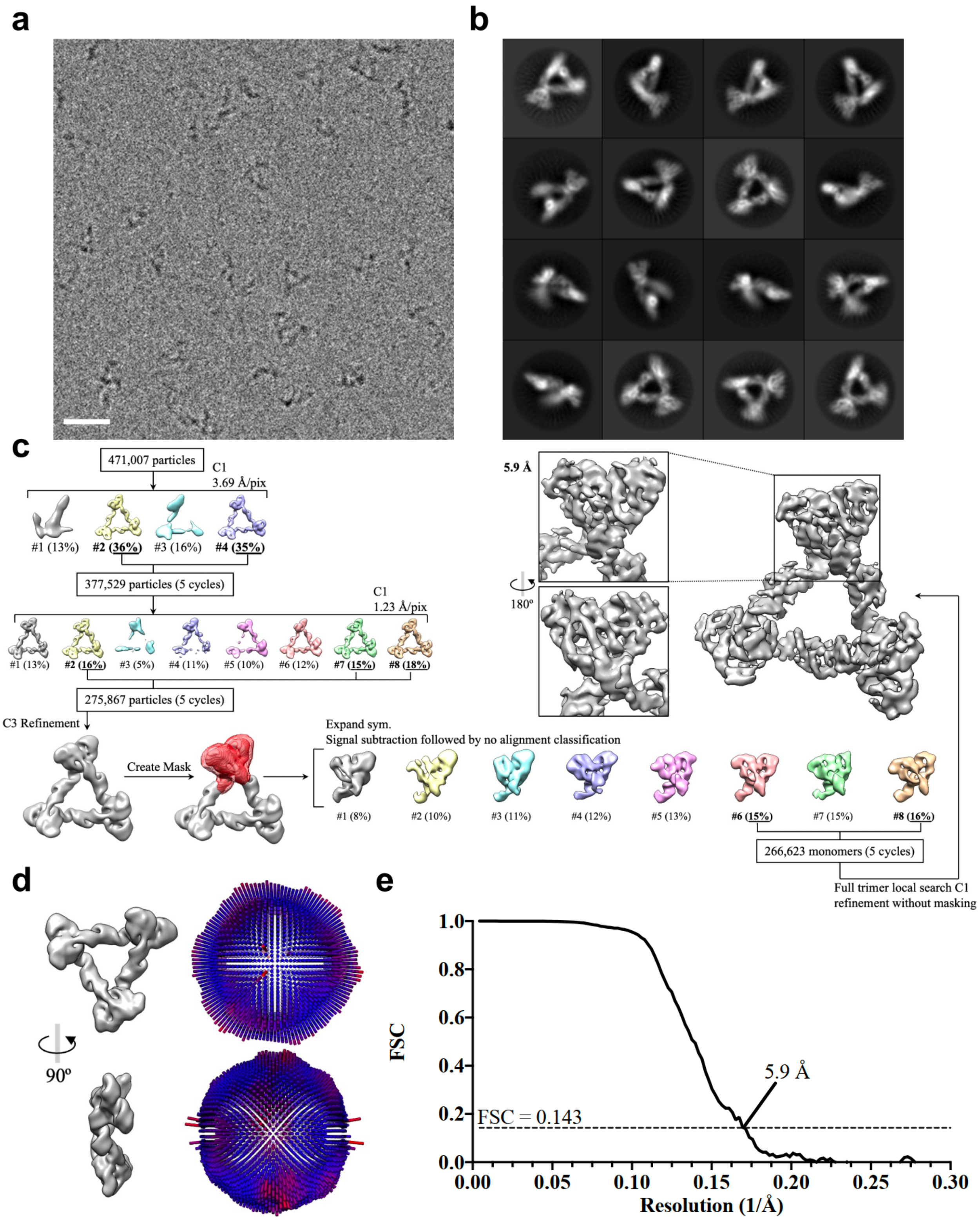


**Supplementary Figure 14 | Cryo-EM imaging, processing and validation for FMNrsw-T. a,** Representative cryo-EM image of FMNrsw-T. The scale bar represents 20 nm. **b,** 2D class averages of FMNrsw-T. Box size is 236 Å. **c,** Processing flowchart for the FMNrsw-T dataset **d,** Angle distribution for the particles included in the final 3D reconstruction. **e,** Gold-standard Fourier Shell Correlation (FSC) for the final FMNrsw-T reconstruction.

| PDB ID | Construct description | Sub-group | Method | Resolution | Active-site metal ions | Reference (Year, Journal) |
|---|---|---|---|---|---|---|
| 1gid | Tet P4-P6 | IC1 | X-ray | 2.50 Å | n.a. | 1996, Science |
| 1hr2 | Tet P4-P6, ΔC209 mutant | IC1 | X-ray | 2.25 Å | n.a. | 2001, Structure |
| 2R8S | Tet P4-P6, ΔC209 mutant, Fab-facilitated crystallization | IC1 | X-ray | 1.95 Å | n.a. | 2007, PNAS |
| 1k2g | Tet P7-P9.0 (G-binding site mimic) | IC1 | NMR | n.a. | n.a. | 2002, RNA |
| 1grz | Tet P3-P9 & P4-P6 (ribozyme core, or Tet3-9) | IC1 | X-ray | 5.0 Å | n.a. | 1998, Science |
| 1x8w | Tet P3-P9 & P4-P6 (ribozyme core, or Tet3-9), 5 mutation sites | IC1 | X-ray | 3.8 Å | 1 (M2) | 2004, Mol. Cell |
| 6wls | Tet ribozyme, truncated at U409, no exon bound | IC1 | cryo-EM (Ribosolve) | 6.8 Å | n.a. | 2020, Nat. Meth. |
| **7r6m** | **Tet post-2SΔP10, dimer with P6b & P8 mutated, 2 deoxy mutations at -1 & +1 residues of ligated exon** | **IC1** | **cryo-EM (ROCK)** | **3.78 (3.68) Å** | **1 (M2)** | **this work** |
| **7r6n** | **Tet ribozyme, trimer with P6b & P9.2 mutated, no exon bound** | **IC1** | **cryo-EM (ROCK)** | **4.17 (4.10) Å** | **n.a.** | **this work** |
| **7r6l** | **Tet pre-2SΔ5'ex, dimer with P6b & P8 mutated, 2 deoxy mutations near scissile phosphate** | **IC1** | **cryo-EM (ROCK)** | **2.98 (2.85) Å** | **1 (M1)** | **this work** |
| 1u6b | Azo Pre-2S, P6a mutated for U1A-facilitated crystallization, 4 deoxy mutations at exon & near scissile phosphate | IC3 | X-ray | 3.10 Å | 2 (monovalent M2) | 2004, Nature |
| 1zzn | Azo Pre-2S, P6a mutated for U1A-facilitated crystallization, 1 deoxy mutation at -1 residue of 5'-exon | IC3 | X-ray | 3.37 Å | 2 | 2005, Science |
| 3bo2 | Azo omit-P, P6a mutated for U1A-facilitated crystallization, the scissile phosphate removed | IC3 | X-ray | 3.31 Å | 2 | 2008, PNAS |
| 3bo4 | Azo post-2S, P6a mutated for U1A-facilitated crystallization, 1 deoxy mutation at -1 residue ligated exon | IC3 | X-ray | 3.33 Å | 2 | 2008, PNAS |
| 3bo3 | Azo pre/post-2S, P6a mutated for U1A-facilitated crystallization, a mixture of two states | IC3 | X-ray | 3.4 Å | 2 | 2008, PNAS |
| **n.a.** | **Azo post-2S, trimer with P5 & P8 mutated, 2 deoxy mutations at -1 & +1 residues of ligated exon** | **IC3** | **cryo-EM (ROCK)** | **4.9 Å** | **n.a.** | **this work** |
| 1y0q | Two ribozyme, mutations at P5a loop for crystallization, 5'-exon bound | IA2 | X-ray | 3.6 Å | 1 (M2) | 2005, Nat. SMB |
| 2rkj | Two ribozyme, 5'-exon bound, co-crystal with CYT-18 protein | IA2 | X-ray | 4.5 Å | n.a. | 2008, Nature |
| 4p8z | DirLC wild-type, a mixture of two states | GIR1 | X-ray | 3.85 Å | n.a. | 2014, PNAS |
| 6gyv | DirLC circularly permutated | GIR1 | X-ray | 2.5 Å | 1 (different reaction) | 2014, PNAS |

**Supplementary Table 1 | Different group I intron structures or fragments that have been determined.** The structures in this work are highlighted in bold. The resolution shown in parentheses are for the TetGI core in this work.

| Sequence | Explanation |
|---|---|
| GGTTCTAATACGACTCACTATA**G**GACCTTTGGAGGGAAAAGTTATCAGGCATGCACCTGGTAGCTAGTCTTTAAACCAATAGATTGCATCGGTTTAAAAGGCAA<br>GACCGTCAAATTGCGGGAAAGGGGTCAACAGCCGTTCAGTACCAAGTCTCAGGGGAAACTTTGAGATGGCCTTGCAAAGGGTATGGTAATAAGCTGACGGACA<br>TGGTCCTAACCACGCAGCCAAGTCCTAAGTCAACAGATCTTCTGTTGATATGGATGCAGTTCACAGACTAAATGTCGGTCGGGGAAGATGTATTCTTCTCATAAG<br>ATATAGTCGGACCTCTCCTTAATGGGAGCTAGCGGATGAAGTGATGCAACACTGGAGCCGCTGGGAACTAATTTGTATGCGAAAGTATATTGATTAGTTTTGGA<br>GTACTC**G***AAGGGCGTCGTCGCCCCGAGCGGTAGTAAGCAGGGAACTCACCTCCAATTTCAGTACTGAAATTGTCGTAGCAGTTGACTACTGTTATGTGATTGGTA*<br>*GAGGCTAAGTGACGGTATTGGCGTAAGTCAGTATTGCAGCACAGCACAAGCCCGCTTGCGAGAAT* | Template for TetGI-M |
| GGTTCTAATACGACTCACTATA**G**GACCTTTGGAGGGAAAAGTTATCAGGCATGCACCTGGTAGCTAGTCTTTAAACCAATAGATTGCATCGGTTTAAAAGGCAA<br>GACCGTCAAATTGCGGGAAAGGGGTCAACAGCCGTTCAGTACCAAGTCTCAGGGGAAACTTTGAGATGGCCTTGCAAAGGGTATGGTAATAAGCTGACGGACA<br>TGGTCCTAACCACGCAGCCAAGTCCTAAGTCAAGGATGGTTCTTGATATGGATGCAGTTCACAGACTAAATGTCGGTCGGGGAAGAGAACCATCCTCTTCTCATA<br>AGATATAGTCGGACCTCTCCTTAATGGGAGCTAGCGGATGAAGTGATGCAACACTGGAGCCGCTGGGAACTAATTTGTATGCGAAAGTATATTGATTAGTTTTG<br>GAGTACTC**G***AAGGGCGTCGTCGCCCCGAGCGGTAGTAAGCAGGGAACTCACCTCCAATTTCAGTACTGAAATTGTCGTAGCAGTTGACTACTGTTATGTGATTGG*<br>*TAGAGGCTAAGTGACGGTATTGGCGTAAGTCAGTATTGCAGCACAGCACAAGCCCGCTTGCGAGAAT* | Template for TetGI-D |
| GGTTCTAATACGACTCACTATA**G**GACCTTTGGAGGGAAAAGTTATCAGGCATGCACCTGGTAGCTAGTCTTTAAACCAATAGATTGCATCGGTTTAAAAGGCAA<br>GACCGTCAAATTGCGGGAAAGGGGTCAACAGCCGTTCAGTACCAAGTCTCAGGGGAAACTTTGAGATGGCCTTGCAAAGGGTATGGTAATAAGCTGACGGACA<br>TGGTCCTAACCACGCAGCCAAGTCCTAAGTCAACAGGATGGTTCTGTTGATATGGATGCAGTTCACAGACTAAATGTCGGTCGGGGAAGATGTATTCTTCTCATA<br>AGATATAGTCGGACCTCTCCTTAATGGGAGCTAGCGGATGAAGTGATGCAACACTGGAGCCGCTGGGAACTAATCAGTGAACCATCCACTGATTAGTTTTGGAG<br>TACTC**G***AAGGGCGTCGTCGCCCCGAGCGGTAGTAAGCAGGGAACTCACCTCCAATTTCAGTACTGAAATTGTCGTAGCAGTTGACTACTGTTATGTGATTGGTAG*<br>*AGGCTAAGTGACGGTATTGGCGTAAGTCAGTATTGCAGCACAGCACAAGCCCGCTTGCGAGAAT* | Template for TetGI-T |
| GGTTCTAATACGACTCACTATA**G**GCCGGGTGGAGGGAAAAGTTATCAGGCATGCACCTGGTAGCTAGTCTTTAAACCAATAGATTGCATCGGTTTAAAAGGCAA<br>GACCGTCAAATTGCGGGAAAGGGGTCAACAGCCGTTCAGTACCAAGTCTCAGGGGAAACTTTGAGATGGCCTTGCAAAGGGTATGGTAATAAGCTGACGGACA<br>TGGTCCTAACCACGCAGCCAAGTCCTAAGTCAACAGATCTTCTGTTGATATGGATGCAGTTCACAGACTAAATGTCGGTCGGGGAAGATGTATTCTTCTCATAAG<br>ATATAGTCGGACCTCTCCTTAATGGGAGCTAGCGGATGAAGTGATGCAACACTGGAGCCGCTGGGAACTAATTTGTATGCGAAAGTATATTGATTAGTTTTGGA<br>GTACTC**G***AAGGGCGTCGTCGCCCCGAGCGGTAGTAAGCAGGGAACTCACCTCCAATTTCAGTACTGAAATTGTCGTAGCAGTTGACTACTGTTATGTGATTGGTA*<br>*GAGGCTAAGTGACGGTATTGGCGTAAGTCAGTATTGCAGCACAGCACAAGCCCGCTTGCGAGAAT* | Template for TetGI G14-G414 |
| GGTTCTAATACGACTCACTATA**G**GCCGGGTGGAGGGAAAAGTTATCAGGCATGCACCTGGTAGCTAGTCTTTAAACCAATAGATTGCATCGGTTTAAAAGGCAA<br>GACCGTCAAATTGCGGGAAAGGGGTCAACAGCCGTTCAGTACCAAGTCTCAGGGGAAACTTTGAGATGGCCTTGCAAAGGGTATGGTAATAAGCTGACGGACA<br>TGGTCCTAACCACGCAGCCAAGTCCTAAGTCAACAGATCTTCTGTTGATATGGATGCAGTTCACAGACTAAATGTCGGTCGGGGAAGATGTATTCTTCTCATAAG<br>ATATAGTCGGACCTCTCCTTAATGGGAGCTAGCGGATGAAGTGATGCAACACTGGAGCCGCTGGGAACTAATTTGTATGCGAAAGTATATTGATTAGTTTTGGA<br>GTACTCGACCCGGCC**A***GGCCGGCATGGTCCCAGCCTCCTCGCTGGCGCCGGCTGGGCAACATGCTTCGGCATGGCGAATGGGAC* | Template for TetGI G14-a+9 |
| GGTTCTAATACGACTCACTATA**G**GCCGGGTGGAGGGAAAAGTTATCAGGCATGCACCTGGTAGCTAGTCTTTAAACCAATAGATTGCATCGGTTTAAAAGGCAA<br>GACCGTCAAATTGCGGGAAAGGGGTCAACAGCCGTTCAGTACCAAGTCTCAGGGGAAACTTTGAGATGGCCTTGCAAAGGGTATGGTAATAAGCTGACGGACA<br>TGGTCCTAACCACGCAGCCAAGTCCTAAGTCAACAGATCTTCTGTTGATATGGATGCAGTTCACAGACTAAATGTCGGTCGGGGAAGATGTATTCTTCTCATAAG<br>ATATAGTCGGACCTCTCCTTAATGGGAGCTAGCGGATGAAGTGATGCAACACTGGAGCCGCTGGGAACTAATTTGTATGCGA**A***GGCCGGCATGGTCCCAGCCTC*<br>*CTCGCTGGCGCCGGCTGGGCAACATGCTTCGGCATGGCGAATGGGAC* | Template for TetGI monomer G14-A386 |
| GGTTCTAATACGACTCACTATA**G**GCCGGGTGGAGGGAAAAGTTATCAGGCATGCACCTGGTAGCTAGTCTTTAAACCAATAGATTGCATCGGTTTAAAAGGCAA<br>GACCGTCAAATTGCGGGAAAGGGGTCAACAGCCGTTCAGTACCAAGTCTCAGGGGAAACTTTGAGATGGCCTTGCAAAGGGTATGGTAATAAGCTGACGGACA<br>TGGTCCTAACCACGCAGCCAAGTCCTAAGTCAAGGATGGTTCTTGATATGGATGCAGTTCACAGACTAAATGTCGGTCGGGGAAGAGAACCATCCTCTTCTCATA<br>AGATATAGTCGGACCTCTCCTTAATGGGAGCTAGCGGATGAAGTGATGCAACACTGGAGCCGCTGGGAACTAATTTGTATGCGA**A***GGCCGGCATGGTCCCAGC*<br>*CTCCTCGCTGGCGCCGGCTGGGCAACATGCTTCGGCATGGCGAATGGGAC* | Template for TetGI dimer G14-A386 (5' IVT RNA of TetGI-DS) |
| GGTTCTAATACGACTCACTATA*GGGCATCAATATACTCTGATGAGTCCGTGAGGACGAAACGAGCTAGCTCGTC***A**GTATATTGATTAGTTTTGGAGTACTCGACC<br>CGGCC**A***GGCCGGCATGGTCCCAGCCTCCTCGCTGGCGCCGGCTGGGCAACATGCTTCGGCATGGCGAATGGGAC* | Template for rTetCIRC |
| GGTTCTAATACGACTCACTATA**G**GCCGTGTGCCTTGCGCCGGGAAACCACGCAAGGGATGGTGTCAAATTCGGCGAAACCTAAGCGCCCGCCCGGGCGTATGG<br>CAACGCCGAGCCAAGCTTCGCAGCCTTCGGGCTGCGATGAAGGTGTAGAGACTAGACGGCACCCACCTAAGGCAAACGCTATGGTGAAGGCATAGTCCAGGG<br>AGTGGCGAAAGCCACACAAACCA**G***AAGGGCGTCGTCGCCCCGAGGATC* | Template for AzoGI-M |
| GGTTCTAATACGACTCACTATA**G**GCCGTGTGCCTTGCGCCGGGAAACCACGCAAGGGATGGTGTCAAATTCGGCGAAACCTAAGCGCGGATGGTTCGCGTATG<br>GCAACGCCGAGCCAAGCTTCGCAGCCTTCGGGCTGCGATGAAGGTGTAGAGACTAGACGGCACCCACCTAAGGCGCACCGAACCATCCGGTGCGCTATGGTGA<br>AGGCATAGTCCAGGGAGTGGCGAAAGCCACACAAACCA**G***AAGGGCGTCGTCGCCCCGAGGATC* | Template for AzoGI-T |
| GGTTCTAATACGACTCACTATA**G**CGGTAGTAAGCAGGGAACTCACCTCCAATTTCAGTACTGAAATTGTCGTAGCAGTTGACTACTGTTATGTGATTGGTAGAGG<br>CTAAGTGACGGTATTGGCGTAAGTCAGTATTGCAGCACAGCACAAGCCCGCTTGCGAGAA**T**GTCCAACCTTCATGCTTACGACG | Template for trans-acting VS ribozyme |
| GGTTCTAATACGACTCACTATA**G**GCCGTGTGCCTTGCGCCGGGAAACCACGCAAGGGATGGTGTCAAATTCGGCGAAACCTAAGCGCCCGCCCGGGCGTATGG<br>CAACGCCGAGCCAAGCTTCGCAGCCTTCGGGCTGCGATGAAGGTGTAGAGACTAGACGGCACCCACCTAAGGCAAACGCTATGGTGAAGGCATAGTCCAGGG<br>AGTGGCGAAAGCCACACAAACCAGACGGCC**A***GGCCGGCATGGTCCCAGCCTCCTCGCTGGCGCCGGCTGGGCAACATGCTTCGGCATGGCGAATGGGAC* | Template for AzoGI*-M |
| GGTTCTAATACGACTCACTATA**G**GCCGTGTGCCTTGCGCCGGGAAACCACGCAAGGGATGGTGTCAAATTCGGCGAAACCTAAGCGCGGATGGTTCGCGTATG<br>GCAACGCCGAGCCAAGCTTCGCAGCCTTCGGGCTGCGATGAAGGTGTAGAGACTAGACGGCACCCACCTAAGGCGCACCGAACCATCCGGTGCGCTATGGTGA<br>AGGCATAGTCCAGGGAGTGGCGAAAGCCACACAAACCAGACGGCC**A***GGCCGGCATGGTCCCAGCCTCCTCGCTGGCGCCGGCTGGGCAACATGCTTCGGCATG*<br>*GCGAATGGGAC* | Template for AzoGI*-T |
| GTTCTAATACGACTCACTATA**G**GATCTTCGGGGCAGGGTGAAATTCCCGACCGGTGGTATAGTCCACGAAAGTATTTGCTTTGATTTGGTGAAATTCCAAAACCG<br>ACAGTAGAGTCTGGATGAGAGAAGATT**C***GGCCGGCATGGTCCCAGCCTCCTCGCTGGCGCCGGCTGGGCAACATGCTTCGGCATGGCGAATGGGAC* | Template for FMNrsw-M |
| GTTCTAATACGACTCACTATA*GGGCAATCAGCGATCCCTGATGAGTCCGTGAGGACGAAACGAGCTAGCTCGTC***G**GATCGCTGATTGATCCGCTTCGGGGCAGG<br>GTGAAATTCCCGACCGGTGGTATAGTCCACGAAAGCGCGAATCAGCCGCGCTTTGATTTGGTGAAATTCCAAAACCGACAGTAGAGTCTGGATGAGAGAAG**C***G*<br>*GCCGGCATGGTCCCAGCCTCCTCGCTGGCGCCGGCTGGGCAACATGCTTCGGCATGGCGAATGGGAC* | Template for FMNrsw-T |

**Supplementary Table 2 | Sequences of synthesized genes for PCR amplification to produce the IVT templates.** The 5'- and 3'ends of the target RNA are in bold. Sequences removed by ribozyme(s) are in italics. Primer-binding regions for PCR amplification are underlined. For the preparation of AzoGI-M and AzoGI-T, the IVT was performed with the trans-acting VS ribozyme, which was introduced by adding a 0.1 equivalent of the PCR-amplified template for trans-acting VS ribozyme.

| | TetGI-DS | TetGI-D | TetGI-T | AzoGI-T | FMNrw-T |
|---|---|---|---|---|---|
| | PDB code: 7R6L EMDB code: EMD-24281 | PDB code: 7R6M EMDB code: EMD-24282 | PDB code: 7R6N EMDB code: EMD-24283 | PDB code: N/A EMDB code: EMD-24284 | PDB code: N/A EMDB code: EMD-24285 |
| **Data collection and processing** | | | | | |
| Microscope | Titan Krios | Titan Krios | Titan Krios | Polara | Polara |
| Detector | K3 | K3 | K3 | K2 Summit | K2 Summit |
| Magnification | 105,000 | 105,000 | 105,000 | 31,000 | 31,000 |
| Voltage (kV) | 300 | 300 | 300 | 300 | 300 |
| Electron Exposure ($e^-/Å^2$) | 47 | 47 | 47 | 52 | 52 |
| Defocus Range (μm) | 0.8-2.0 | 0.8-2.0 | 0.8-2.0 | 1.0-2.5 | 1.0-2.5 |
| Acquisition Pixel Size (Å) | 0.825 | 0.825 | 0.825 | 1.23 | 1.23 |
| Final symmetry imposed | C1 | C1 | C1 | C1 | C1 |
| Initial particle images (#) | 550,754 | 384,170 | 124,353 | 474,349 | 471,007 |
| Final particle images (#) | 82,575 S.E. monomers | 113,548 S.E. monomers | 85,596 S.E. monomers | 486,860 S.E. monomers | 266,623 S.E. monomers |
| Reconstruction Pixel Size (Å) | 0.825 | 1.03 | 1.24 | 1.23 | 1.23 |
| Map resolution (Å) | 2.85 core / 2.98 overall | 3.68 core / 3.78 overall | 4.10 core / 4.17 overall | 4.9 | 5.9 |
| FSC threshold | 0.143 | 0.143 | 0.143 | 0.143 | 0.143 |
| Map resolution range (Å) | 2.5-6.0 | 3.5-8.0 | 3.0-7.0 | 4.0-8.0 | 4.0-8.0 |
| Refinement | | | | | |
| Model resolution (Å) | 2.97 | 3.82 | 4.20 | | |
| FSC threshold | 0.5 | 0.5 | 0.5 | | |
| Map sharpening B factor ($Å^2$) | N/A | N/A | N/A | N/A | N/A |
| Model composition | | | | | |
| Non-hydrogen atoms | 7857 | 7755 | 7588 | | |
| RNA bases | 366 | 362 | 354 | | |
| Ligand ($Mg^{2+}$ ions) | 20 | 13 | 12 | | |
| B factor ($Å^2$) | | | | | |
| RNA bases | 53.79 | 100.66 | 89.86 | | |
| Ligand ($Mg^{2+}$ ions) | 19.61 | 55.48 | 46.19 | | |
| R.m.s. deviations | | | | | |
| Bond lengths (Å) | 0.008 | 0.006 | 0.008 | | |
| Bond angles (°) | 0.893 | 0.943 | 1.069 | | |

**Supplementary Table 3 | Data collection and refinement statistics.**

**Supplementary references**


1. Bindewald, E., Grunewald, C., Boyle, B., O'Connor, M. & Shapiro, B. A. Computational strategies for the automated design of RNA nanoscale structures from building blocks using NanoTiler. *J Mol Graph Model* 2008, **27**(3)**:** 299-308.
2. Tomizawa, J.-i. Control of ColE1 plasmid replication: The process of binding of RNA I to the primer transcript. *Cell* 1984, **38**(3)**:** 861-870.
3. Lee, A. J. & Crothers, D. M. The solution structure of an RNA loop–loop complex: the ColE1 inverted loop sequence. *Structure* 1998, **6**(8)**:** 993-1007.
4. Guo, F., Gooding, A. R. & Cech, T. R. Structure of the Tetrahymena ribozyme: base triple sandwich and metal ion at the active site. *Mol Cell* 2004, **16**(3)**:** 351-362.
5. Lehnert, V., Jaeger, L., Michel, F. & Westhof, E. New loop-loop tertiary interactions in self-splicing introns of subgroup IC and ID: a complete 3D model of the Tetrahymena thermophila ribozyme. *Chemistry & Biology* 1996, **3**(12)**:** 993-1009.
6. Adams, P. L., Stahley, M. R., Kosek, A. B., Wang, J. & Strobel, S. A. Crystal structure of a self-splicing group I intron with both exons. *Nature* 2004, **430**(6995)**:** 45-50.
7. Serganov, A., Huang, L. & Patel, D. J. Coenzyme recognition and gene regulation by a flavin mononucleotide riboswitch. *Nature* 2009, **458**(7235)**:** 233-237.
8. Zuker, M. Mfold web server for nucleic acid folding and hybridization prediction. *Nucleic Acids Res.* 2003, **31**(13)**:** 3406-3415.
9. Liu, D. *et al.* Branched kissing loops for the construction of diverse RNA homooligomeric nanostructures. *Nature Chem.* 2020, **12**(3)**:** 249-259.
10. Denesyuk, N. A. & Thirumalai, D. How do metal ions direct ribozyme folding? *Nat Chem* 2015, **7**(10)**:** 793-801.
11. Schorb, M., Haberbosch, I., Hagen, W. J. H., Schwab, Y. & Mastronarde, D. N. Software tools for automated transmission electron microscopy. *Nat Methods* 2019, **16**(6)**:** 471-477.
12. Zheng, S. Q. *et al.* MotionCor2: anisotropic correction of beam-induced motion for improved cryo-electron microscopy. *Nat Methods* 2017, **14**(4)**:** 331-332.
13. Rohou, A. & Grigorieff, N. CTFFIND4: Fast and accurate defocus estimation from electron micrographs. *J Struct Biol* 2015, **192**(2)**:** 216-221.
14. Shaikh, T. R. *et al.* SPIDER image processing for single-particle reconstruction of biological macromolecules from electron micrographs. *Nat Protoc* 2008, **3**(12)**:** 1941-1974.
15. Zivanov, J. *et al.* New tools for automated high-resolution cryo-EM structure determination in RELION-3. *Elife* 2018, **7**.
16. Scheres, S. H. Processing of Structurally Heterogeneous Cryo-EM Data in RELION. *Methods Enzymol.* 2016, **579:** 125-157.
17. Kucukelbir, A., Sigworth, F. J. & Tagare, H. D. Quantifying the local resolution of cryo-EM density maps. *Nat Methods* 2014, **11**(1)**:** 63-65.
18. Terwilliger, T. C., Ludtke, S. J., Read, R. J., Adams, P. D. & Afonine, P. V. Improvement of cryo-EM maps by density modification. *Nat Methods* 2020, **17**(9)**:** 923-927.
19. Adams, P. D. *et al.* PHENIX: a comprehensive Python-based system for macromolecular structure solution. *Acta Crystallogr D Biol Crystallogr* 2010, **66**(Pt 2)**:** 213-221.
20. Pettersen, E. F. *et al.* UCSF Chimera--a visualization system for exploratory research and analysis. *J Comput Chem* 2004, **25**(13)**:** 1605-1612.
21. Afonine, P. V. *et al.* Real-space refinement in PHENIX for cryo-EM and crystallography. *Acta Crystallogr D Struct Biol* 2018, **74**(Pt 6)**:** 531-544.
22. Emsley, P. & Cowtan, K. Coot: model-building tools for molecular graphics. *Acta Crystallogr D Biol Crystallogr* 2004, **60**(Pt 12 Pt 1)**:** 2126-2132.
23. McGreevy, R., Teo, I., Singharoy, A. & Schulten, K. Advances in the molecular dynamics flexible fitting method for cryo-EM modeling. *Methods* 2016, **100:** 50-60.
24. Johnston, W. K., Unrau, P. J., Lawrence, M. S., Glasner, M. E. & Bartel, D. P. RNA-catalyzed RNA

polymerization: accurate and general RNA-templated primer extension. *Science* 2001, **292**(5520)**:** 1319-1325.

25. Zaug, A. J., Kent, J. R. & Cech, T. R. A labile phosphodiester bond at the ligation junction in a circular intervening sequence RNA. *Science* 1984, **224**(4649)**:** 574-578.
26. Zhang, K. *et al.* Cryo-EM structure of a 40 kDa SAM-IV riboswitch RNA at 3.7 A resolution. *Nat Commun* 2019, **10**(1)**:** 5511.
27. Wang, J. & Moore, P. B. On the interpretation of electron microscopic maps of biological macromolecules. *Protein Sci.* 2017, **26**(1)**:** 122-129.
28. Cate, J. H. *et al.* Crystal structure of a group I ribozyme domain: principles of RNA packing. *Science* 1996, **273**(5282)**:** 1678-1685.
29. Cate, J. H., Hanna, R. L. & Doudna, J. A. A magnesium ion core at the heart of a ribozyme domain. *Nat Struct Biol* 1997, **4**(7)**:** 553-558.
30. Ye, J. D. *et al.* Synthetic antibodies for specific recognition and crystallization of structured RNA. *Proc. Natl. Acad. Sci. USA* 2008, **105**(1)**:** 82-87.
31. Pyle, A. M. & Cech, T. R. Ribozyme recognition of RNA by tertiary interactions with specific ribose 2'-OH groups. *Nature* 1991, **350**(6319)**:** 628-631.
32. Herschlag, D., Eckstein, F. & Cech, T. R. Contributions of 2'-hydroxyl groups of the RNA substrate to binding and catalysis by the Tetrahymena ribozyme. An energetic picture of an active site composed of RNA. *Biochemistry* 1993, **32**(32)**:** 8299-8311.
33. Strobel, S. A. & Cech, T. R. Tertiary interactions with the internal guide sequence mediate docking of the P1 helix into the catalytic core of the Tetrahymena ribozyme. *Biochemistry* 1993, **32**(49)**:** 13593-13604.
34. Golden, B. L., Kim, H. & Chase, E. Crystal structure of a phage Twort group I ribozyme-product complex. *Nat Struct Mol Biol* 2005, **12**(1)**:** 82-89.
35. Suh, S. O., Jones, K. G. & Blackwell, M. A Group I intron in the nuclear small subunit rRNA gene of Cryptendoxyla hypophloia, an ascomycetous fungus: evidence for a new major class of Group I introns. *J. Mol. Evol.* 1999, **48**(5)**:** 493-500.
36. Leontis, N. B. & Westhof, E. A common motif organizes the structure of multi-helix loops in 16 S and 23 S ribosomal RNAs. *J Mol Biol* 1998, **283**(3)**:** 571-583.
